# Transcriptional profiling of three *Pseudomonas syringae* pv. *actinidiae* biovars reveals different responses to apoplast-like conditions related to strain virulence

**DOI:** 10.1101/2020.08.11.246074

**Authors:** Elodie Vandelle, Teresa Colombo, Alice Regaiolo, Tommaso Libardi, Vanessa Maurizio, Davide Danzi, Annalisa Polverari

## Abstract

*Pseudomonas syringae* pv. *actinidiae* (Psa) is a phytopathogen that causes devastating bacterial canker in kiwifruit. Among five biovars defined by genetic, biochemical and virulence traits, Psa3 is the most aggressive and is responsible for the most recent reported outbreaks, but the molecular basis of its heightened virulence is unclear. We therefore designed the first *P. syringae* multi-strain whole-genome microarray, encompassing biovars Psa1, Psa2 and Psa3 and the well-established model *P. syringae* pv. *tomato*, and analyzed early bacterial responses to an apoplast-like minimal medium. Transcriptomic profiling revealed (i) the strong activation in Psa3 of all *hrp*/*hrc* cluster genes, encoding components of the type III secretion system required for bacterial pathogenicity and involved in responses to environmental signals; (ii) potential repression of the *hrp*/*hrc* cluster in Psa2; and (iii) activation of flagellum-dependent cell motility and chemotaxis genes in Psa1. The detailed investigation of three gene families encoding upstream regulatory proteins (histidine kinases, their cognate response regulators, and proteins with diguanylate cyclase and/or phosphodiesterase domains) indicated that c-di-GMP may be a key regulator of virulence in Psa biovars. The gene expression data were supported by the quantification of biofilm formation. Our findings suggest that diverse early responses to the host apoplast, even among bacteria belonging to the same pathovar, can lead to different virulence strategies and may explain the differing outcomes of infections. Based on our detailed structural analysis of *hrp* operons, we also propose a revision of *hrp* cluster organization and operon regulation in *P. syringae.*

**Author summary:** *Pseudomonas syringae* pv. *actinidiae* (Psa) is a bacterial pathogen that infects kiwifruit crops. Recent outbreaks have been particularly devastating due to the emergence of a new biovar (Psa3), but the molecular basis of its virulence is unknown so it is difficult to develop mitigation strategies. In this study, we compared the gene expression profiles of Psa3 and various less-virulent biovars in an environment that mimics early infection, to determine the basis of pathogenicity. Genes involved in the assembly and activity of the type III secretion system, which is crucial for the secretion of virulence effectors, were strongly upregulated in Psa3 while lower or not expressed in the other biovars. We also observed the Psa3-specific expression of genes encoding upstream signaling components, confirming that strains of the same bacterial pathovar can respond differently to early contact with their host. Finally, our data suggested a key role in Psa virulence switch ability for the small chemical signaling molecule c-di-GMP, which suppresses the expression of virulence genes. This effect of c-di-GMP levels on Psa3 virulence should be further investigated and confirmed to develop new mitigation methods to target this pathway.

## Introduction

*Pseudomonas syringae* pv. *actinidiae* (Psa) is a Gram-negative pathogenic bacterium that causes bacterial canker in kiwifruit. In the last 10 years, severe outbreaks of the disease worldwide have caused huge economic losses, particularly in Italy, New Zealand and China, which are among the largest producers. Following an epiphytic phase of multiplication, Psa can enter kiwifruit plants via natural openings such as stomata and hydathodes, by mechanical wounding, or by pollen dissemination (1). Symptoms include blossom necrosis and brown–black leaf spots often surrounded by a chlorotic halo. From primary infection sites, bacteria can move systemically through the twigs and trunk, generating extensive cankers that produce white-to-orange exudates (2).

Five different biovars of Psa have been defined based on genetic, biochemical and biological traits, including virulence and toxin production. Biovar 1 (here described as Psa1), which can synthesize phaseolotoxin, includes the original Psa strains found in Japan in 1984. Biovar 2 (here described as Psa2) has only been found in Korea (1992–1997), and produces the toxin coronatine. Biovar 3 (here described as Psa3) was responsible for the severe outbreaks of bacterial canker in Italy, New Zealand, Chile and China between 2008 and 2010, and is the most virulent despite the apparent absence of toxins (3). Biovar 4, identified in Australia and New Zealand, was initially classified as a low-virulence Psa strain but was subsequently reclassified as a different pathovar: *Pseudomonas syringae* pv. *actinidifoliorum* (4)(5). Finally, biovars 5 and 6 were found in small areas in Japan in 2012 and 2015, respectively, and are poorly characterized (6,7). Given the major differences in biovar behavior and the severe economic losses associated with recent Psa3 outbreaks, it is increasingly important to gain insight into the evolution and pathogenicity of Psa, and this has elevated the species to the status of a model organism for the analysis of bacterial infections in woody plants. In particular, we need to understand the virulence of the most aggressive strains and the triggers that allow endophytic Psa populations to become virulent, which could facilitate the development of new control strategies.

Effector proteins are essential bacterial pathogenicity factors exported into plant cells by the specialized type III secretion system (TTSS). They subvert host cellular processes to support bacterial proliferation, thus promoting disease. TTSS components are encoded by hypersensitive reaction and pathogenicity (*hrp*) and *hrp* conserved (*hrc*) genes (8–10), which are present in almost all phytopathogenic bacteria including Psa (11,12). The expression of *hrp/hrc* genes is one of the most important events during infection, and is induced following contact with plant tissues or in minimal *hrp*-inducing medium (HIM) (13–16). The latter is widely used to study the behavior of bacteria in contact with host cells because it reproduces apoplast-like conditions, including minimal levels of nutrients and glucose/fructose as available carbon sources, and a low pH (15).

As a first step in the search for Psa3 virulence determinants, genomic analysis revealed genes encoding four TTSS-related effector proteins (HopH1, HopZ5, HopAM1-2 and HopAA1-2) that are missing in other biovars, and may therefore account for the aggressiveness of Psa3 (11,12). However, this is not sufficient evidence to confirm the molecular basis of differential virulence because it is not clear whether or not these genes are expressed. Transcriptomic analysis is therefore required to determine gene expression profiles under different conditions, revealing how the bacteria adapt to interactions with host cells or new growth conditions, as well as shedding light on the regulatory networks that govern such adaptive responses. HIM is often used in this context to trigger the expression of TTSS genes (15).

To identify Psa genes and pathways associated with the activation of virulence and strain-dependent aggressiveness, we designed a custom multi-strain microarray covering the pan-genome of Psa1, Psa2 and Psa3 as well as the model organism *P. syringae* pv. *tomato* DC3000 (Pto). We carried out a comprehensive transcriptomic analysis of these biovars to identify differentially expressed genes following the inoculation of minimal medium (HIM) compared to King’s B (KB) rich medium, the latter representing standard *in vitro* growth conditions. We also tested two different Psa3 strains representing the two recent outbreaks that occurred simultaneously in New Zealand (ICMP18884/V-13) and Italy (CRAFRU8.43).

Hypotheses based on the transcriptomic data were explored using a bioinformatics pipeline to compare (1) the structural features of specific Psa genomic regions, and (2) the whole-genome identification and transcriptomic characterization of three bacterial gene families acting as virulence activators or TTSS regulators: histidine kinases, their response regulators, and proteins involved in c-di-GMP metabolism and signaling. The results highlighted structural and functional differences that might account for the differential aggressiveness and diverse virulence mechanisms in the three Psa biovars, and also questioned widely-accepted paradigms of bacterial pathogenicity by indicating the ability of even biovars within the same *P. syringae* pathovar to deploy different responses and pathogenicity mechanisms following contact with the host. Our study therefore provides new insight into the molecular basis of virulence in bacterial pathogens that infect tree crops.

## Results

### Multi-strain microarray design

To find transcriptomic differences among different Psa biovars, we designed a multi-strain microarray containing the whole set of annotated sequences from four Psa strains belonging to the three main biovars best characterized at the time: NCPPB3739 (Psa1), ICMP19073 (Psa2), ICMP18884 (Psa3) and CRA-FRU8.43 (Psa3). The Pto DC3000 genome was included as a control to distinguish general and Psa-specific transcriptomic features. Finally, we also included annotated transcripts from three integrative conjugative element (ICE) sequences. Annotated transcripts on the multi-strain microarray (representing all *P. syringae* strains and ICE sequences) were collected as cDNA sequences from different public sources (**Table S1**) in July 2015. Additional genome annotations representing the CRA-FRU 8.43 (11) strain were obtained from the University of Udine (Prof. Giuseppe Firrao, personal communication).

We carried out *in silico* comparisons to identify sub-lists of shared and strain-specific sequences from the cDNA libraries allowing us to prepare a non-redundant cDNA collection comprising the unique multi-strain set of annotated protein coding sequences. These BLAST comparisons identified a subset of genes shared by all strains (*core genome*) as well as subsets of partially shared and strain-specific genes (*dispensable genome*) The multi-strain protein sequence dataset (20,561 sequences) was uploaded to the Agilent eArray web-based application for the design of a five-genome *P. syringae* high-density microarray for single-strain and cross-strain comparative transcriptomics (**Figure S1**). Extended datasets for the sequences and bioinformatics pipeline described herein can be found in **Appendices S1**-**S7**.

### Assignment of proteins to orthologous clusters for cross-strain comparison

To define groups of orthologous sequences, we used reciprocal best hits (17–21) to detect orthologous genes in our *Pseudomonas* dataset based on significant reciprocal similarity between amino acidic sequences. We used BLASTP (22) to perform all pairwise sequence alignments between any two proteins encoded in the five *Pseudomonas* genomes and three ICE sequences, and recorded the best hit for each protein.

Close cross-strain investigation of the strain-specific and multi-strain sets of protein coding genes highlighted residual redundancy that failed to capture obvious relationships due to minimal differences in the annotated protein sequences. To remove this confounding redundancy and improve the performance of the microarray in cross-strain gene expression comparisons, we further grouped the proteins into super-clusters based on slightly relaxed values in the output of BLASTP all-against-all runs (E < 0.001; at least 94% identity over at least 90% of the aligned sequences). The above procedure assigned the full and redundant collection of protein sequences collected for different *P. syringae* strains from different annotation sources (**Fig S1**) to a non-redundant set of 10,839 protein groups (**S7 Appendix**).

The identified protein groups were then compared across *P. syringae* strains to identify sets of protein-coding genes shared by all strains (*core genome*) as well as partially shared and strain-specific genes (*dispensable genome*) (23). This analysis defined a core Psa/Pto genome of 3551 genes, which expanded to 4206 genes when we only considered the sequences common to Psa biovars (**Fig 1A**). This broadly agreed with a previous report showing that Psa includes 3916 single-copy orthologs shared by four biovars: Psa1–3 and Psa5 (24). Finally, the comparison of strains CRAFRU8.43 and ICMP18884 revealed a core Psa3 genome of 6103 unique supercluster IDs and strain-specific accessory genomes of 307 and 73 unique IDs, respectively (**Fig 1B**). This broadly agrees with the 363 clade-specific genes previously identified in Psa3 strains (24).

**Fig 1.**
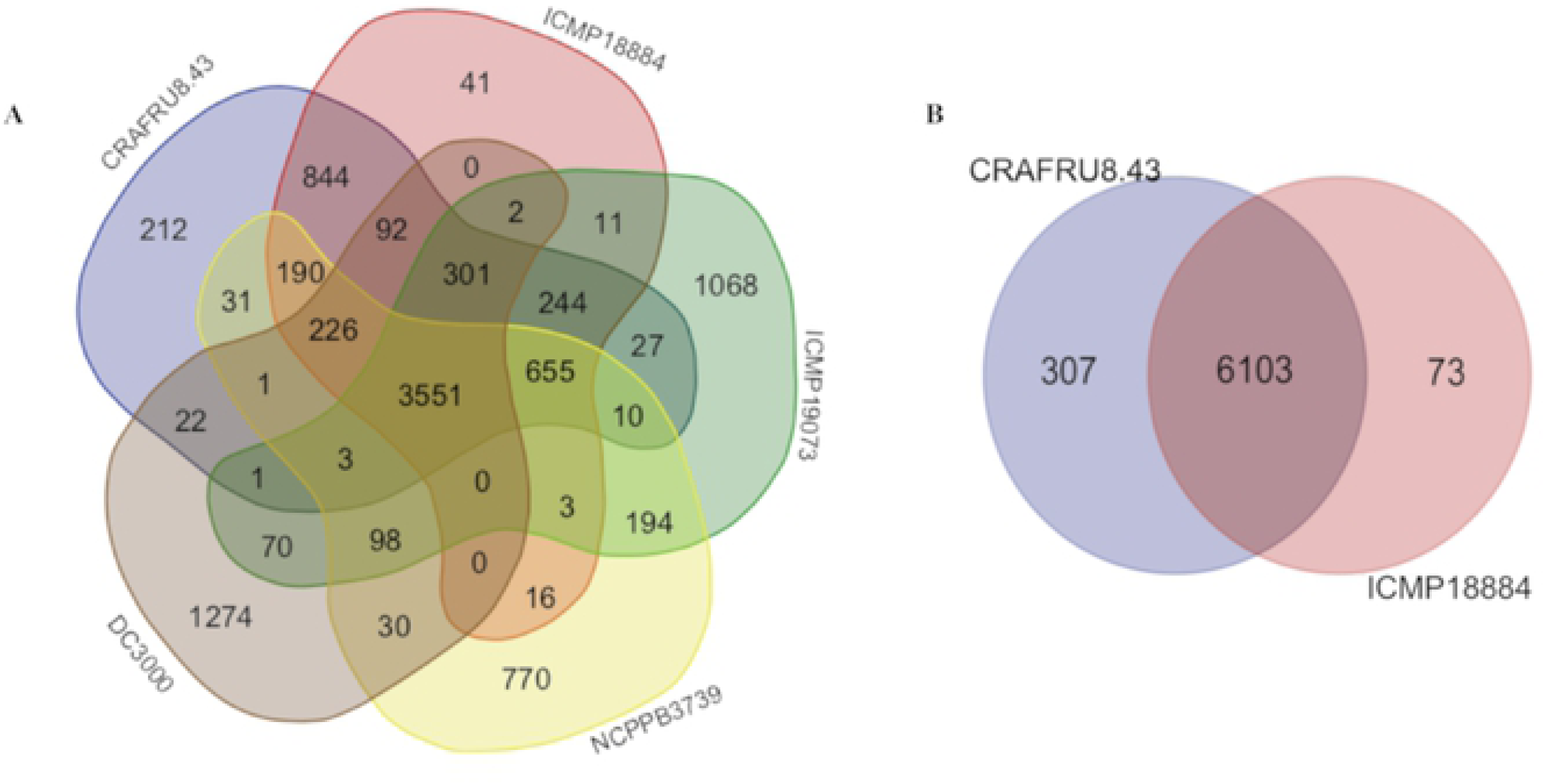
*Core* and *dispensable* proteomes of the Psa strains. The Venn diagrams show the common and unique proteins in (A) the five Psa strains considered in this study and (B) in the strains CRAFRU8.43 and ICMP18884 representing Psa3. The Venn diagrams were generated using Draw Venn Diagram (http://bioinformatics.psb.ugent.be/webtools/Venn/) based on superclusters to avoid protein redundancy among strains.

### General transcriptional profiles of Psa biovars and Pto under apoplast-like conditions

Next, the four Psa strain were cultured in rich medium (KB) or minimal medium (HIM) to identify genes modulated specifically under apoplast-like conditions. Samples were harvested after 4 or 8 h of incubation for microarray analysis. The hierarchical clustering of samples based on the similarity of gene expression profiles resulted in dendrograms with high reproducibility among the biological replicates (**Fig 2**). The first level of clustering showed a clear separation between the pathovars Psa and Pto, whereas the second level separated the Psa3 strains (CRAFRU8.43 and ICMP18884) from Psa1 (J35) and Psa2 (KN.2). Finally, the third level separated Psa1 and Psa2. The growth conditions (rich or minimal medium) and sampling time points (4 or 8 h) played only a minor role in the clustering of Psa strains, but the separation of rich and minimal medium was clearly observed for Pto. Principal component analysis (PCA) confirmed the clustering revealed by the dendrogram (**Fig S2**). Overall, these results indicated that microarray data extrapolated from the experiment can identify genes with statistically significant differential expression among different pathovars or biovars and different media.

**Fig 2.**
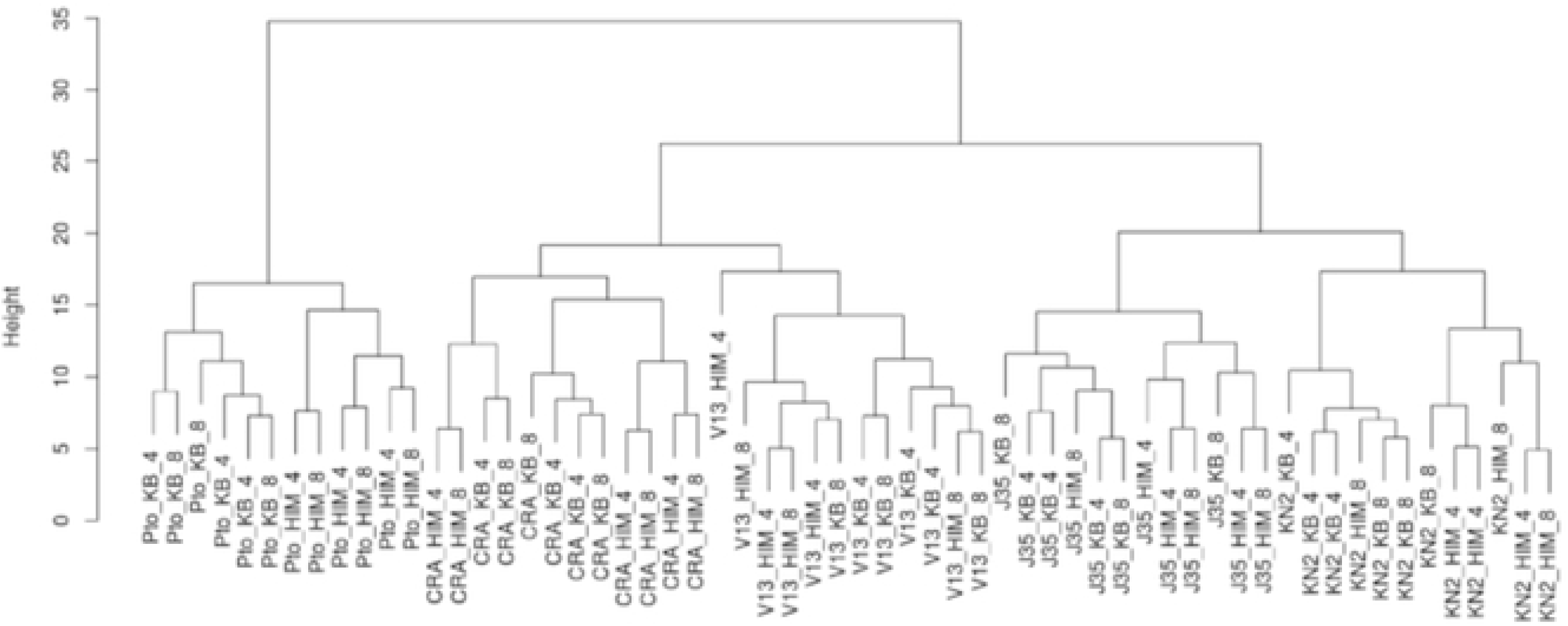
Clustering of samples based on gene expression profiles as determined by microarray analysis. Each sample is represented by a composite identifier, in which the first field shows the strain (possible values: Pto = Pto DC3000, CRA= CRAFRU 8.43, V13 = ICMP18884, J35 = NCPPB3739/ICMP9617, KN2 = KN.2), the second represents the growth conditions (possible values: KB = King’s B medium, HIM = *hrp*-inducing medium), and the third represents the harvest time point (possible values: 4 or, 8 h post-inoculation).

Preliminary analysis revealed a smaller number of differentially expressed genes (DEGs) in the Psa3 strain CRAFRU8.43 (914 DEGs) than in the others (∼1500 DEGs) when comparing growth on minimal and rich medium at both time points, with a false discovery rate (FDR) < 0.05 and an absolute log_2_ fold change (log_2_FC) > |1| (**Fig 3A; S1-S10 Datasets**). The removal of genes modulated in the same manner at both time points confirmed this trend, revealing 648 DEGs in CRAFRU8.43 compared to ∼1000–1400 DEGs for the others (**Fig S3**). Overall, ∼50% of genes were upregulated and ∼50% were downregulated in all strains. When considering the time points individually, the two Psa3 strains (CRAFRU8.43 and ICMP18884) behaved in a similar manner, with most DEGs detected at 8 h, whereas Psa1, Psa2 and Pto featured a higher number of DEGs at 4 h. Although this indicates that Psa3 responds slowly to apoplast-like conditions and shows a later response peak than the other strains, more detailed analysis (described below) indicated that a rapid response to nutrient depletion in the Psa3 strains may have masked the modulation of gene expression at 4 h. Finally, a comparison of DEGs between 4 and 8 h for each strain individually revealed that ∼30% of DEGs were commonly modulated at both time points in Psa3 (CRAFRU8.43 and ICMP18884) as well as Pto, whereas this proportion dropped to 10–15% for Psa1 and Psa2, suggesting that these latter strains respond to minimal medium by regulating different metabolic processes progressively over time (**Fig 3B**).

**Fig 3.**
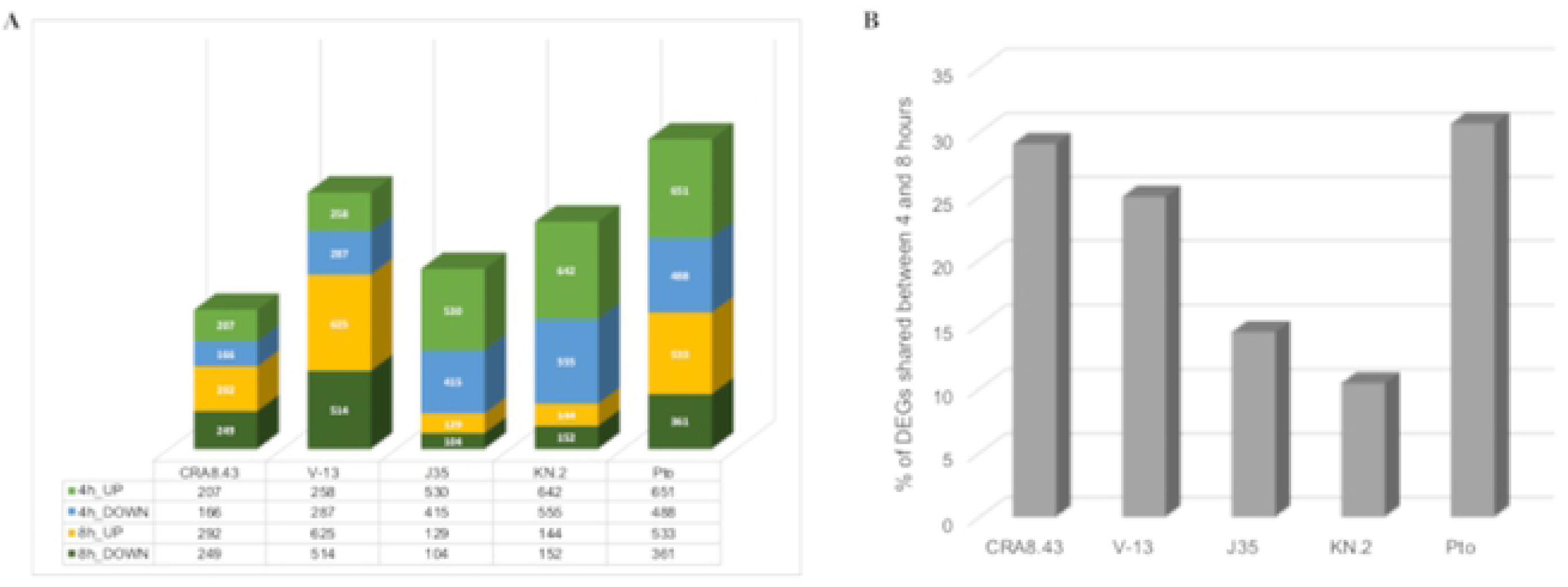
Transcriptome profiles of different Psa strains grown under apoplast-like conditions. (A) Number of differentially expressed genes (DEGs) in the different Psa strains incubated for 4 or 8 h in *hrp*-inducing minimal medium compared with KB rich medium. (B) Number of DEGs commonly regulated at both time points in the different Psa strains expressed as a percentage of the total number of DEGs in each strain.

### Comparison of DEGs among bacterial strains

Next we compared transcriptional data at both time points (without redundancy) to avoid any ‘time effect’ on gene expression. As stated above, strain CRAFRU8.43 appeared less responsive to minimal medium than the other strains. The comparison of CRAFRU8.43 and ICMP18884 showed that almost 80% of DEGs in CRAFRU8.43 (495 of 648) were also modulated in ICMP18884, suggesting a similar response to growth conditions but involving fewer genes (**Fig S4**). To focus on the differences between Psa biovars, we therefore grouped the two Psa3 strains for further analysis.

Genes induced in minimal medium were compared between strains to identify common and unique upregulated genes. The Venn diagram in **Fig 4A** shows that 150 genes were commonly induced in all strains, including genes encoding products related to starvation (e.g., a glycerol uptake facilitator, the phosphate starvation-inducible protein PslF, and members of the sodium:solute symporter protein family) (**Dataset S11**). Furthermore, 56 of these genes corresponded to hypothetical proteins, the identification of which may provide further insights into the response of *P. syringae* to apoplast-like conditions. Finally, we observed the common strong upregulation (log_2_FC > 5) of a gene encoding a C1 family papain-like cysteine protease. Only 51 genes were induced in all Psa strains but not in Pto, suggesting there was no clear pathovar-specific reaction to minimal medium (**Fig 4A**; **Dataset S11**). There were many similarities between Psa3 and Pto, with 126 commonly-induced genes not regulated in Psa1 and Psa2, including 41 encoding TTSS components and others encoding flagellar components or enzymes involved in pyoverdine synthesis, which are important aspects of virulence (**Dataset S11**). Unique genes in both strains also encoded TTSS components, whereas additional Pto-specific genes were involved in coronatine synthesis and Psa-specific genes encoded several peptidases. This suggests that Pto and Psa3 share some common mechanisms for the induction of virulence but also trigger specific virulence pathways. Whereas most strains featured 250–300 uniquely upregulated genes (representing ∼30% of the induced genes in each strain), only 89 such genes were found in Psa1, suggesting a less-specific response (**Dataset S11**). These genes encoded several proteins related to c-di-GMP signaling or chemotaxis. In Psa2, genes related to classical virulence mechanisms were not upregulated, but the unique genes included many related to malonate or sarcosine metabolism, as well as sugar transport (**Dataset S11**). Finally, 92 Psa2 genes corresponded to hypothetical proteins, the function of which will provide insight into the behavior of this strain in the apoplast. Interestingly, although the Psa2 genome contains genes for the complete coronatine biosynthesis pathway, none of these genes was induced under our experimental conditions.

**Fig 4.**
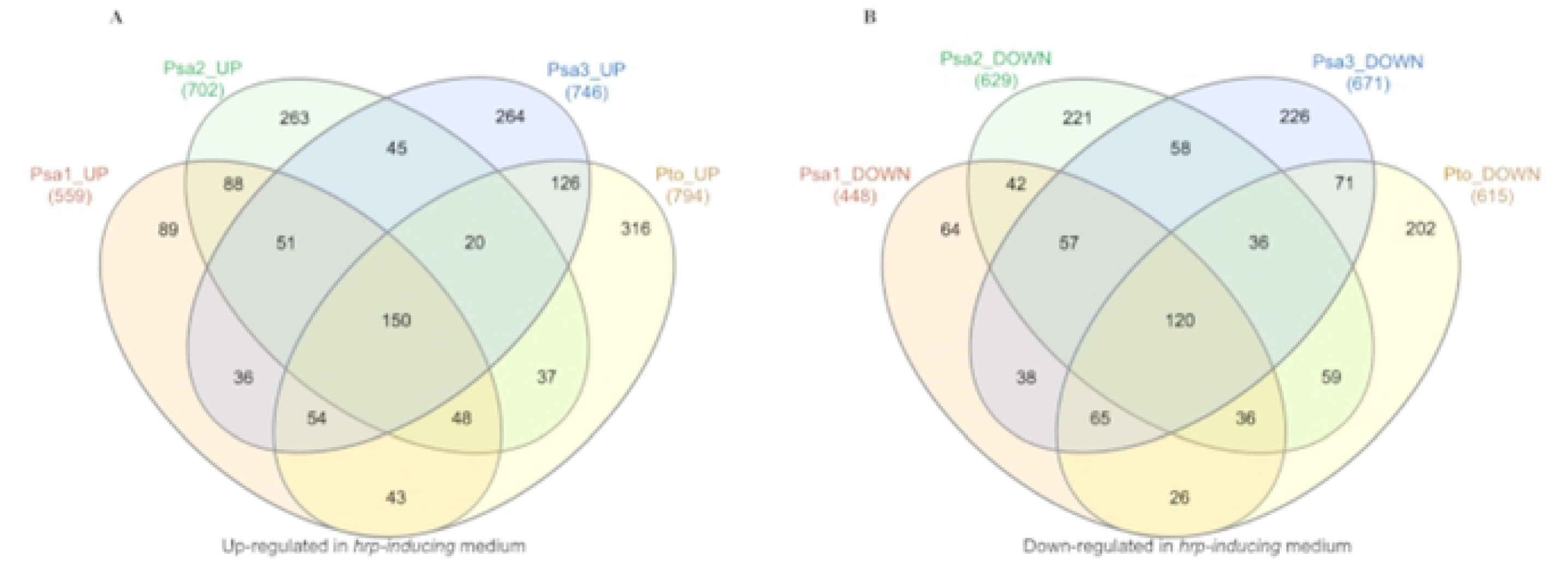
Comparison of genes modulated in *hrp*-inducing medium in different Psa strains. The Venn diagrams show common and unique DEGs (A) upregulated or (B) downregulated in the different strains. The Venn diagrams were generated using Draw Venn Diagram (http://bioinformatics.psb.ugent.be/webtools/Venn/).

Genes suppressed in minimal medium were compared between strains to identify common and unique downregulated genes (**S12 Dataset**). The Venn diagram in **Fig 4B** revealed a similar distribution to the upregulated genes discussed above. We identified 120 genes repressed in all strains, including many encoding ribosomal proteins or with other roles in translation. Only 57 genes were repressed specifically in Psa, with no enrichment for particular functions. Among the 71 genes commonly downregulated in Psa3 and Pto, we observed the gene encoding Lon protease, a negative regulator of the TTSS, consistent with the upregulation of TTSS-related genes in these strains. In most strains, we detected ∼200 uniquely downregulated genes. Again the exception was Psa1, where the number dropped to 64, supporting the lower specificity of its response.

### Biovar-specific responses to apoplast-like conditions

To identify the pathways potentially triggered in each strain by the perception of minimal medium, we used GO analysis to reveal the functionally enriched categories among the genes upregulated in minimal medium (**Fig 5**). Only three categories were commonly enriched in all strains (transcription, styrene catabolism, and vesicle organization) whereas the most enriched categories in Psa3 and Pto (defined as –log_10_(1/adj. p-value) ≥ 15) were related to pathogenesis, the TTSS, and flagellum-dependent cell motility. Another three categories were solely enriched in Psa3 (iron transmembrane transport, glycerol-3-phosphate transmembrane transport, and fumarate metabolism), whereas sporulation and organic hydroxyl compound transport functions were solely enriched in Pto. In contrast, the GO terms that were most enriched in Psa1 included categories related to flagellum-dependent cell motility, chemotaxis and signal transduction, whereas less-enriched categories related to polyhydroxybutyrate biosynthesis and disaccharide metabolism were solely enriched in Psa1. GO enrichment analysis in Psa2 revealed an overall lower degree of enrichment compared to other strains (–log_10_(1/adj. p-value = 3–4) and highlighted functional categories related to signal transduction and the metabolism of carbohydrates, alditol phosphate, branched amino acids (valine and leucine), thioester, cyanate, xylulose, acetyl-CoA and olefin.

**Fig 5.**
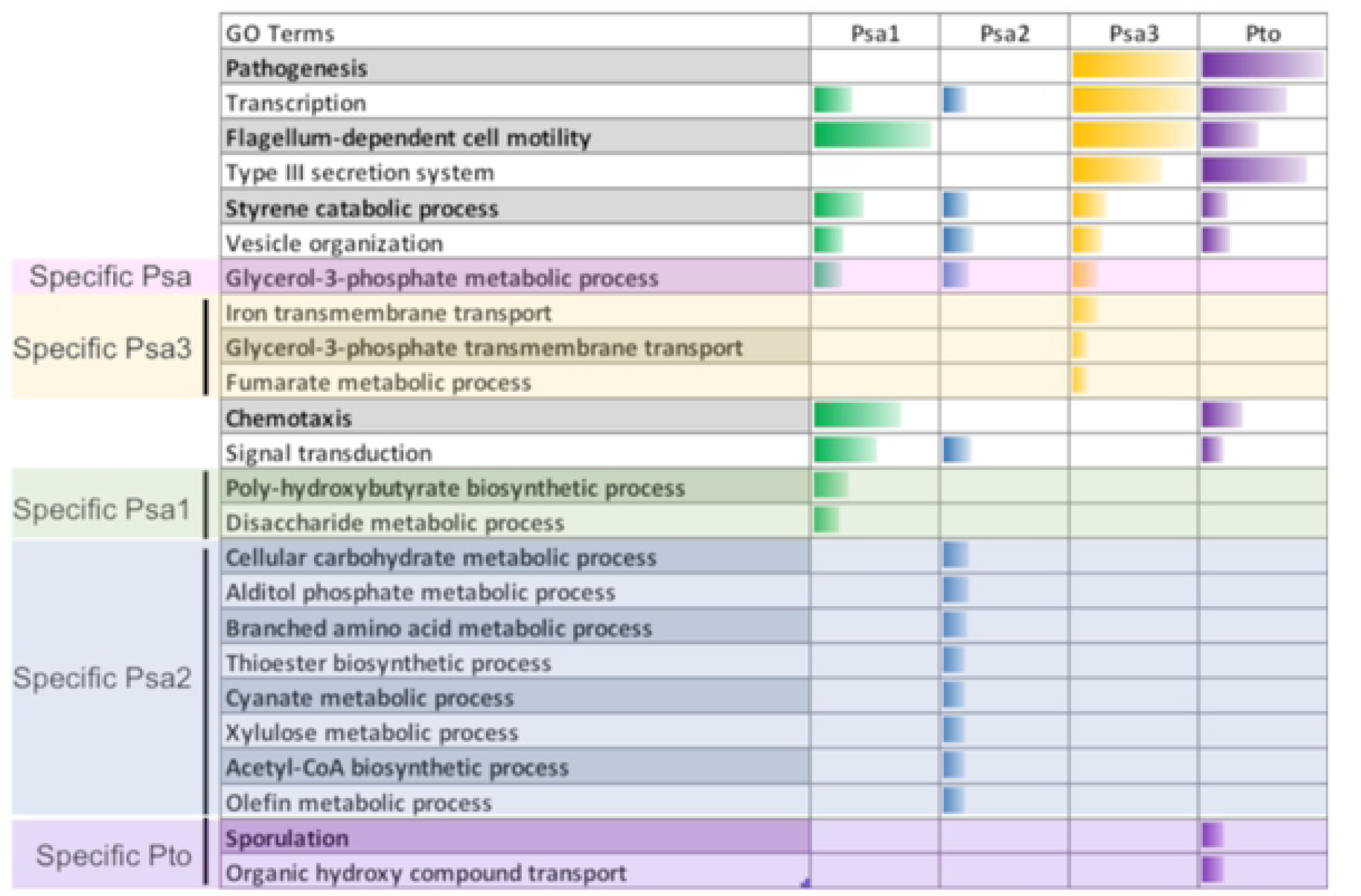
Gene Ontology categories enriched among upregulated genes in different Psa strains under apoplast-like conditions. The data show clusters of GO categories overrepresented in DEGs that are upregulated in different Psa strains in *hrp*-inducing minimal medium compared to rich KB medium, relative to the whole genome. Cluster enrichment was determined using Blast2GO Pro (https://www.blast2go.com).

When we applied the same strategy to identify enriched GO categories among the genes downregulated in minimal medium, the functions of the most enriched categories in each strain (**Fig 6**) and the functions that were enriched solely in a particular strain (**Fig 7**) were more diverse, pointing to a profound reprogramming of global metabolism. The downregulated genes in all strains were enriched for functions related to translation, ATP synthesis, mitochondrial transport, and glycine betaine biosynthesis, suggesting the general reallocation of energy reserves (**Fig 6A**). Some categories were uniquely enriched among the downregulated genes in Psa, including those related to xanthine metabolism and ribosome biogenesis (**Fig 6B**). Others were shared between Psa3 and Pto, including functions related to ion homeostasis, citrate transport, lysine, proline and gamma-aminobutyric acid catabolism, folic acid biosynthesis, and heat responses (**Fig 6C**). Several enriched GO categories were shared by at least two strains (**Fig 6D**). GO categories that were enriched among the downregulated genes uniquely in a given strain included the removal of superoxide radicals in Psa1; branched-amino acid transport, carbohydrate utilization and sucrose metabolism, lipid A and lipopolysaccharide core region biosynthesis, and aromatic amino acid biosynthesis in Psa2; and bacteriocin immunity, guanosine-containing compound biosynthesis, and superoxide metabolism Psa3 (**Fig 7**).

**Fig 6.**
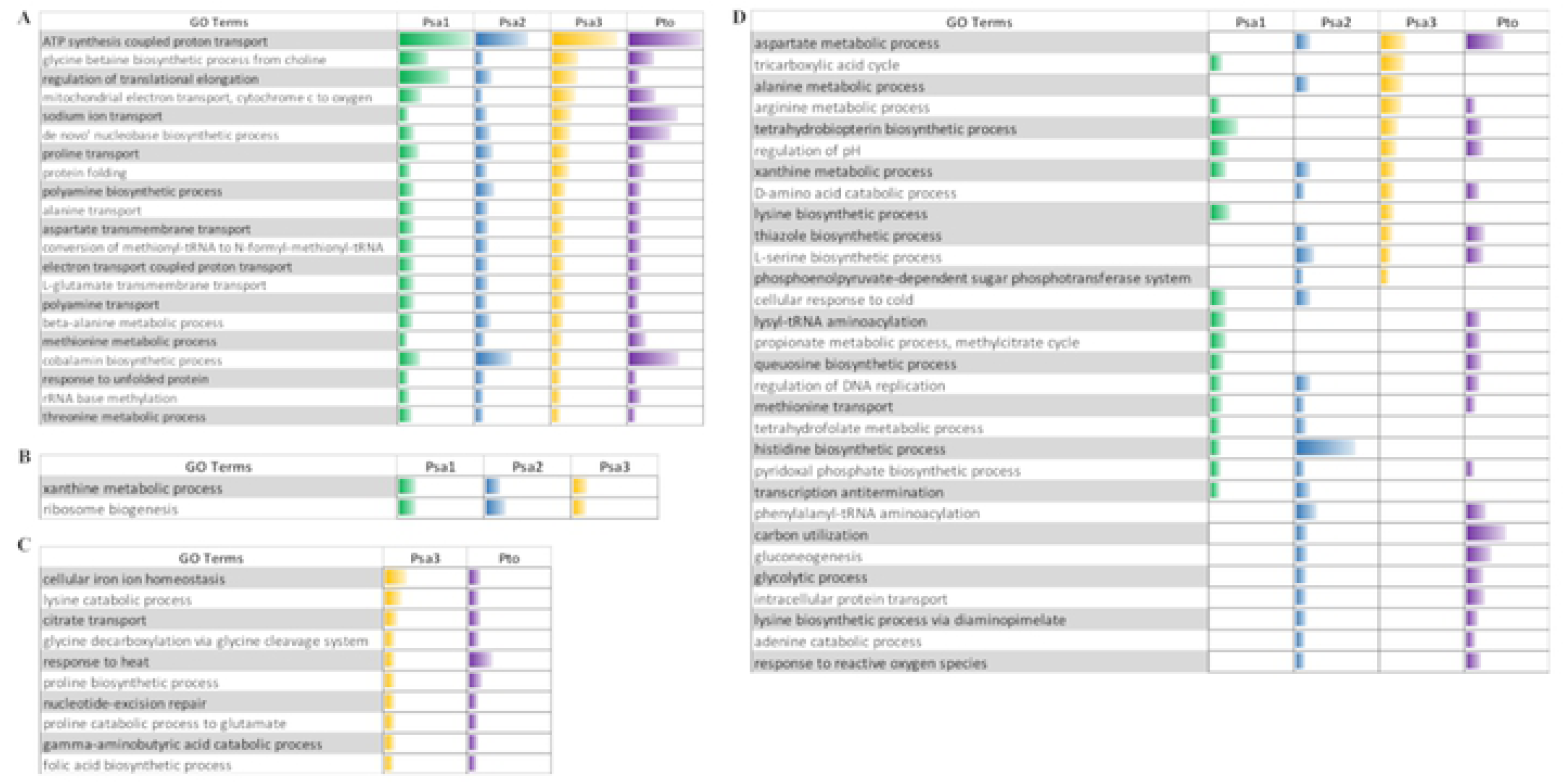
Gene Ontology categories enriched among downregulated genes in different Psa strains under apoplast-like conditions. The data show clusters of GO categories overrepresented in DEGs that are downregulated in different Psa strains. (A) DEGs common to all Psa strains. (B) DEGs unique to single Psa strains. (C) DEGs shared only by Psa3 and Pto. (D) DEGs shared by at least two Psa strains. Cluster enrichment was determined using Blast2GO Pro (https://www.blast2go.com).

**Fig 7.**
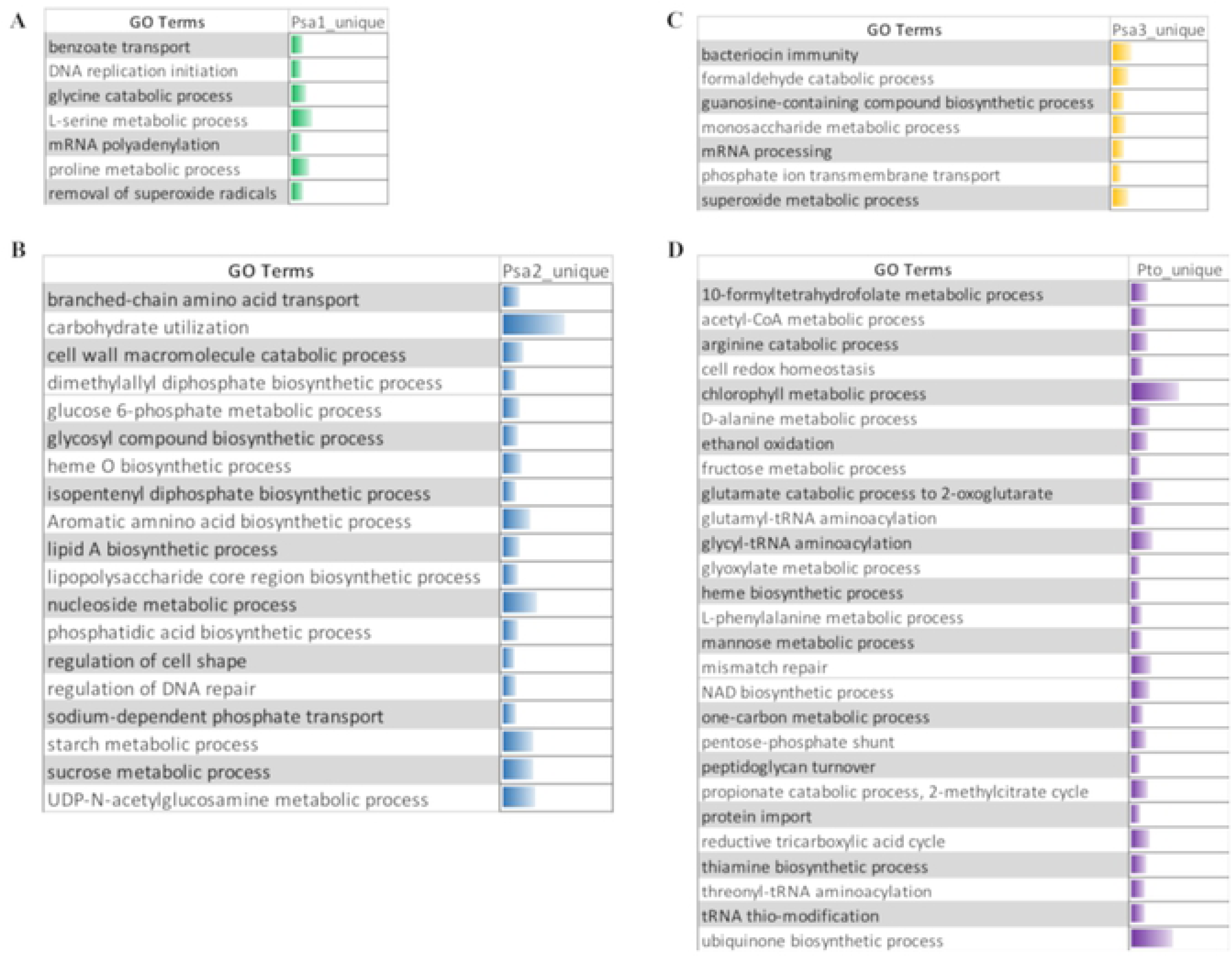
Gene Ontology categories among downregulated genes enriched in specific Psa strains grown under apoplast-like conditions. The data show clusters of GO categories overrepresented in DEGs that are downregulated in specific Psa strains: (A) Psa1, (B) Psa2, (C) Psa3, and (D) Pto. Cluster enrichment was determined using Blast2GO Pro (https://www.blast2go.com).

Interestingly, sample clustering based on enriched GO categories among the upregulated genes indicated that Psa3 and Pto shared similar traits, with a clear induction of pathogenesis-related processes, in contrast to Psa1 and Psa2 (**Fig S5A**). However, sample clustering based on enriched GO categories among downregulated genes showed greater similarity between Psa1 and Psa3 on one hand, and Psa2 and Pto on the other (**Fig S5B**). This confirms that transcriptome profiling is needed to fully understand the similarities and differences among bacterial strains, and indicates that the behavior of Psa1 is intermediate between the mild Psa2 and virulent Psa3 strains.

Another interesting observation concerned the *hrp*/*hrc* cluster, which contains 26 open reading frames organized in seven operons (8). The analysis of gene expression within the *hrp*/*hrc* cluster (**Fig 8A**) showed that all genes, except those encoding the upstream regulators HrpR and HrpS, were strongly upregulated in Psa3 and also in Pto (**Fig S6**). In contrast, only a few genes (*hrpA1, hrpZ, hrpF* and *hrpO*) were induced in Psa1, and the expression level was lower than in Psa3 (log_2_FC ∼4 in Psa3 and ∼2 in Psa1). Only *hrpT* was induced in Psa2, in common with the other strains.

**Fig 8.**
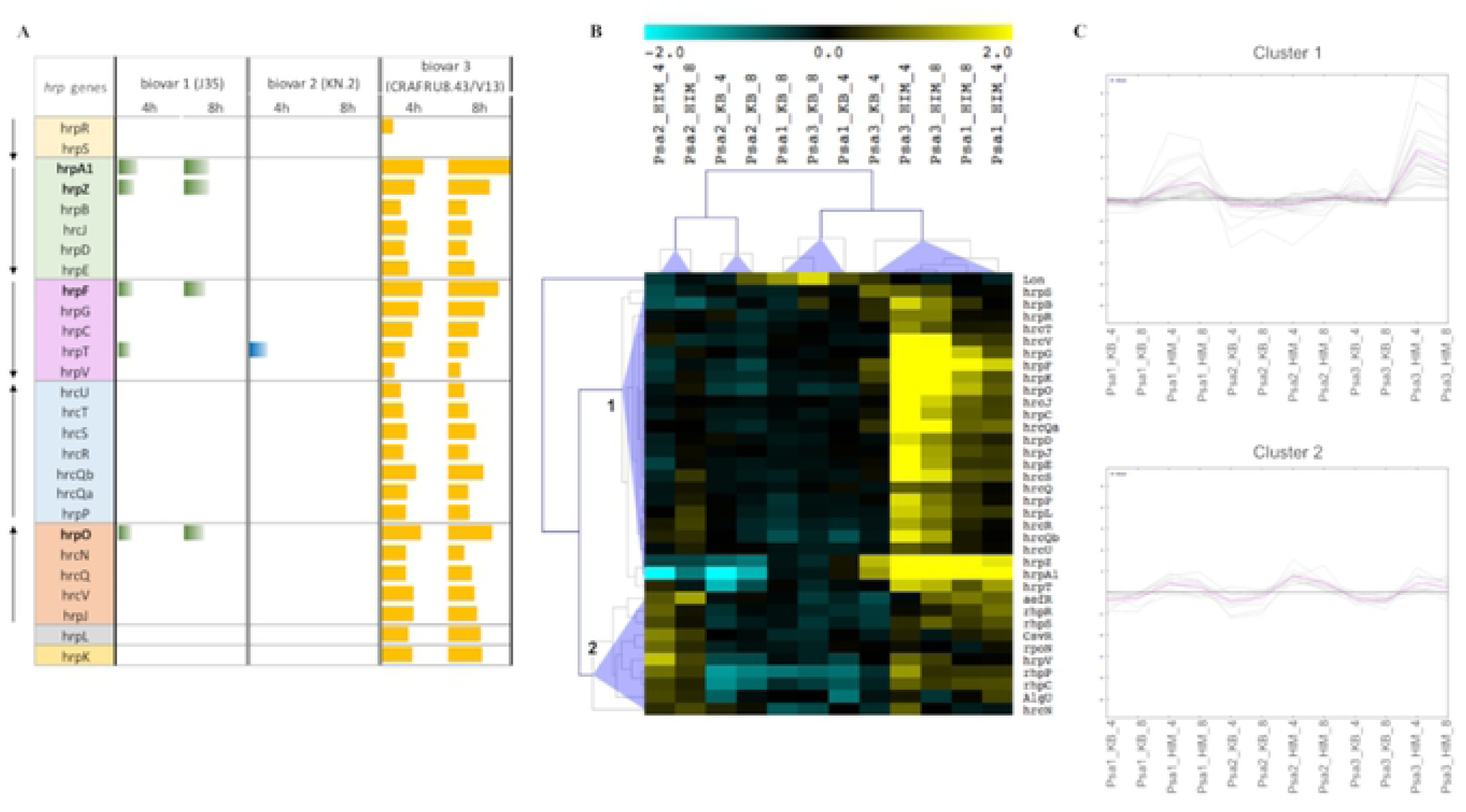
Modulation of *hrp*/*hrc* genes and TTSS regulators in Psa strains incubated in minimal medium. (A) Graphical representation of *hrp*/*hrc* gene expression in the different Psa strains. Each bar represents the log_2_ fold change of expression for the corresponding gene. Color blocks indicate the organization of *hrp*/*hrc* genes in operons. Arrows on the left indicate operon orientation. (B) Hierarchical clustering of the absolute expression level of the *hrp*/*hrc* genes and the main TTSS regulators in the different Psa strains in minimal (HIM) and rich (KB) medium at 4 and 8 h. The clusters were generated using MeV with normalized fluorescence values. Blue triangles indicate the different clusters. (C) Graphical representation of the trend in gene expression for clusters 1 and 2 in the different Psa strains.

To confirm the differential response of the TTSS in the Psa biovars, we transformed them with a reporter system in which the expression of green fluorescent protein (GFP) is driven by the *hrpA1* promoter (p*hrpA1*-GFP). The modified strains were used to monitor *hrpA1* promoter activation as a marker of TTSS activity in minimal medium (**Fig S7**). In agreement with the microarray data, *hrpA1* promoter activity was strong in Psa3, already significant after incubation for 1 h in minimal medium, whereas the induction was slower and ∼50% weaker in Psa1, and no signal was detected in Psa2 over a period of 6 h, confirming that the TTSS is not induced in this strain under these conditions.

### Biovar-dependent responses of the TTSS

Given the importance of the TTSS in bacterial pathogenicity, and based on the differential expression of *hrp*/*hrc* genes in the Psa biovars described above, we investigated the correlation between *hrp*/*hrc* gene expression and virulence in more detail. Hierarchical clustering of *hrp*/*hrc* gene expression profiles across all samples using microarray normalized fluorescence values, including genes encoding the major regulators AefR, RhpR/RhpS, RhpC/RhpP, RpoN, AlgU, CsvR/CsvS and Lon (25), revealed two main clusters. The first included all modulated *hrp*/*hrc* genes except *hrpT, hrpV* and *hrcN* whereas the second included the major TTSS regulators (**Fig 8B**). In cluster 1, Psa3 clearly responded more strongly to minimal medium compared to rich medium, and a similar but weaker response was also observed for Psa1, whereas Psa2 showed no significant modulation in response to minimal medium. In contrast, all Psa biovars showed identical responses to minimal medium in the second cluster at both time points (**Fig 8C**). Surprisingly, all the positive regulators of TTSS were significantly induced in Psa2 after only 4 h of incubation in minimal medium, whereas only some of them were induced in Psa1 and Psa3 (**Fig S8**). In contrast, the negative regulator Lon protease was repressed solely in Psa3. The same hierarchical clustering approach showed that the expression level of *hrp*/*hrc* genes and TTSS regulators was higher in Psa3 grown in rich medium at 4 h compared to other rich medium samples but declined at 8 h. This indicates that at least some *hrp*/*hrc* genes and regulators are already induced by the overnight incubation of Psa3 in rich medium prior to the switch to minimal medium, potentially explaining the apparent weaker response of Psa3 at 4 h.

### The conserved structure of the *hrp*/*hrc* cluster in different Psa biovars

The *hrp*/*hrc* cluster expression profiles revealed that the genes significantly upregulated in Psa1 almost invariably corresponded to the first genes of each operon, based on the previously described operon structure (8), with *hrpO* as the only exception. This was mirrored by the expression profiles in Psa3, where the abovementioned genes were expressed more strongly than the others. These data support the general observation that gene expression in operons declines with distance from the transcriptional start, due to the progressively increasing likelihood of ribosomal detachment (26). Although the general proximal–distal expression gradient supports the reliability of our microarray data, *hrpO* displayed a higher log_2_FC value than the other genes in the *hrpJ* operon (*hrcN, hrcQ, hrcV* and *hrpJ*) and the *hrp* box associated with the *hrpP* gene is > 300 bp upstream, not as close to the transcriptional start site as other *hrp* boxes in the cluster (**Fig S9**). The role of the *hrp* box in the regulation of *hrpP* expression is therefore unclear. We propose a hypothesis in which *hrpO* is not the last gene of the *hrpJ* operon as currently assumed, but the first gene of the *hrpU* operon which lies immediately downstream. We first tested our hypothesis *in silico* by operon prediction using the Psa ICMP18884 genome as a template, which suggested the presence of a single operon spanning all the genes from *hrpJ* to *hrcU* (**Fig S10**). Given these inconclusive data, we tested for the presence of polycistronic transcripts in the Psa3 strain CRAFRU8.43. First, we detected the presence of individual transcripts for *hrcN, hrpO* and *hrpP*, and their expression only in response to minimal medium confirmed our microarray data (**Fig S11**). We then designed primers that spanned pairs of genes (*hrcN*-*hrpO* and *hrpO*-*hrpP*) and detected amplicons only when using the *hrpO*-*hrpP* primers, thus demonstrating that *hrpO* and *hrpP* produce a polycistronic transcript and are part of a single operon (**Fig 9A**). This confirms *hrpO* is the first gene of the *hrpU* operon and lies just upstream of *hrpP*, and further implies that the *hrp* box previously assumed to regulate *hrpP* may actually regulate *hrpO*, even though it is found within the *hrpO* coding sequence (**Fig 9B**). The lack of expression in Psa1 and Psa2 made it impossible to test for polycistronic transcripts in these biovars, but the same operon structure was confirmed by the presence of polycistronic transcripts in Pto (**Fig S12**). These data suggest that the organization of operons *hrpJ* and *hrcU* may be conserved among *P. syringae* strains and that the presence of an upstream *hrp* box does not necessarily specify the positon of an operon.

**Fig 9.**
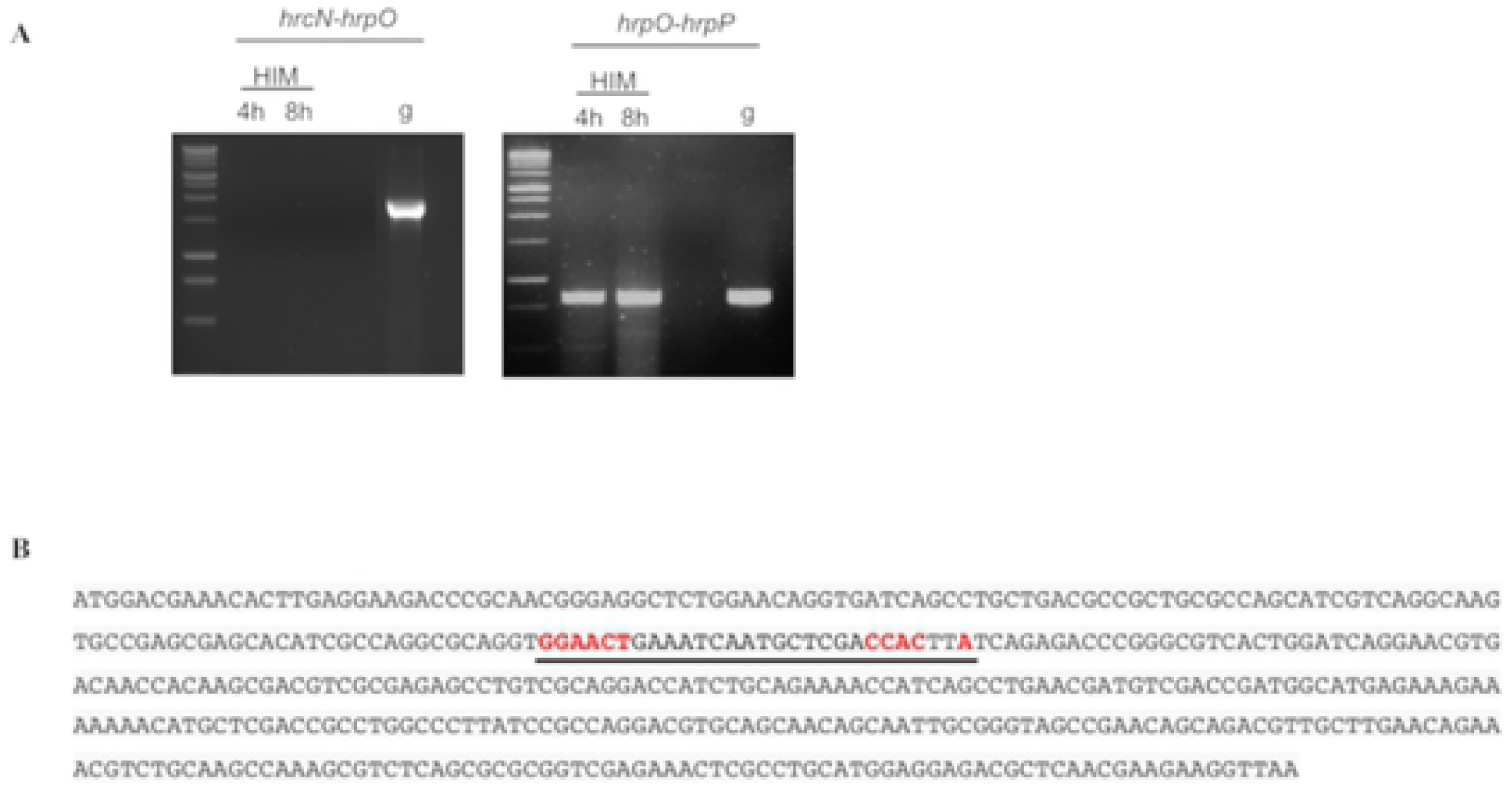
Proposed revision of *hrpJ* and *hrcU* operon organization in Psa3. (A) RT-PCR analysis of polycistronic transcripts containing the *hrpN, hrpO* and *hrpP* sequences in Psa3 (CRAFRU8.43). Primers were designed to amplify polycistronic transcripts encoded by the *hrpN*-*hrpO* and *hrpO*-*hrpP* genes. Total RNA extracted from CRAFRU8.43 cells cultured for 4 or 8 h in minimal medium (HIM) was reverse transcribed and amplified by PCR. Genomic DNA (g) was used as positive control. (B) Coding sequence of *hrpO* including the *hrp* box (underlined). The nucleotides highlighted in red correspond to the *hrp* box consensus motif (GGACC-N_15_-CCAC-N_2_-A).

The complete sequences of the *hrp*/*hrc* clusters from each Psa biovar were aligned to identify genomic differences that might explain the strong induction of *hrp*/*hrc* gene expression in Psa3, the weak induction in Psa1, and absence of a response in Psa2. The *hrp*/*hrc* clusters of Psa1 and Psa3 were closely related, with few nucleotide substitutions, whereas weaker similarity was observed between Psa1/Psa3 and Psa2, in particular in the region covering the *hrcU* operon (**Fig 10A**). To ensure that sequence variations in the *hrcU* operon did not reflect poor genome annotation, we further aligned the *hrp*/*hrc* clusters of two additional Psa2 genomes (ICMP19071 and ICMP19072, which showed high identity with ICMP19073), confirming that the sequence of the Psa2 *hrcU* operon diverges from the other two biovars (**Fig 10B**). However, the nucleotide variations between Psa2 and Psa3 translated to 98–100% identity at the protein level, suggesting most of the nucleotide changes resulted in synonymous codons. Overall, these data indicate that the modulation of *hrp*/*hrc* cluster genes in different Psa biovars is probably related to different signaling pathways and regulators rather than differences in the *hrp*/*hrc* genome sequence.

**Fig 10.**
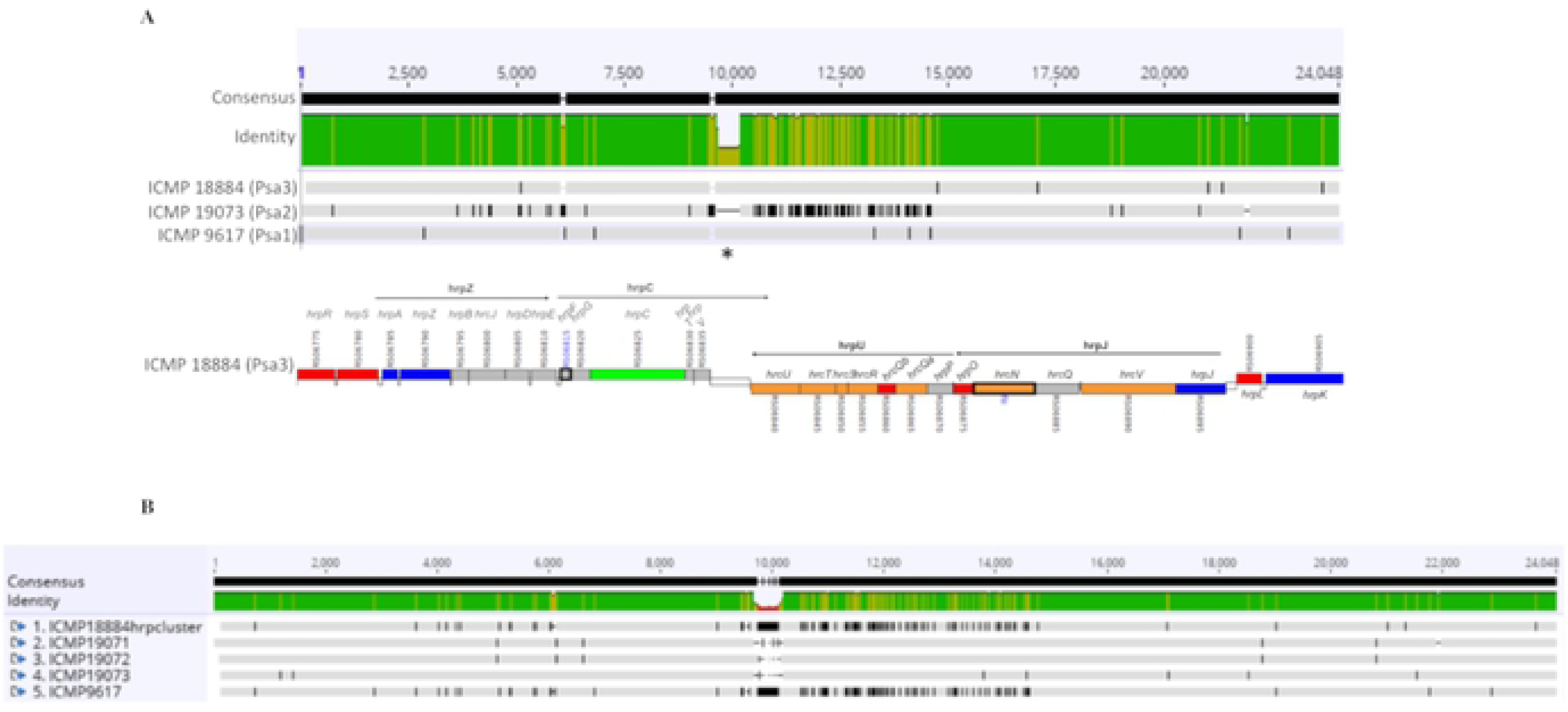
Alignment of the *hrp* clusters from Psa1, Psa2 and Psa3 and cluster organization. (A) Multiple alignment of *hrp* cluster genomic sequences (*hrpR* to *hrpK*) retrieved for three strains of Psa representative of the three major biovars: Psa1 (ICMP9617), Psa2 (ICMP19073) and Psa3 (ICMP18884). Black lines indicate polymorphisms. The lower part shows a graphical representation of *hrp* cluster organization in ICMP18884 with the 27 *hrp*/*hrc* genes and the corresponding operons (8). (B) Multiple alignment of *hrp* cluster genomic sequences from ICMP18884 (Psa3), ICMP9617 (Psa1) and three strains representative of Psa2 (ICMP19071, ICMP19072, ICMP19073). Black lines indicate polymorphisms. Genome sequences were retrieved from the Pseudomonas Genome Database and multiple alignments were prepared using Geneious v11.0.3 (http://www.geneious.com).

### Common and biovar-specific expression profiles of key signaling proteins

In bacteria, signal transduction is largely mediated by members of the histidine kinase (HK) family, which phosphorylate downstream response regulator (RR) proteins as part of a two-component system (27). In some cases, the HK and RR domains are present on the same hybrid histidine kinase (HHK) protein (28). We therefore focused on the role of these key signaling proteins in the response of Psa biovars to the apoplast-like conditions imposed by minimal medium.

Putative HKs were identified using MIST3, retrieving 65, 61 and 66 proteins containing at least one putative HK domain in the genomes of Psa1, Psa2 and Psa3, respectively. Additional candidates were identified by using BLASTP to screen all HKs from each biovar against the genomes of the other two biovars, and by screening all three biovars with the HKs identified in Pto (29). Finally, a whole-genome BLASTN search using probes corresponding to HK sequences confirmed, at the nucleotide level, the presence or absence of further putative HKs. Overall, the analysis identified 72 common candidates (21 HHKs and 51 HKs) in all Psa strains, as well as three (two HHKs and one HK) unique to Psa2, although the two HHKs were also represented in the Psa3 genome as pseudogenes (**Fig 11**). We found that 29 of the genes were differentially expressed (24 induced and five repressed), including 16 that were modulated in Psa1 (15 induced and one repressed), 22 that were modulated in Psa2 (17 induced and five repressed), and only 11 that were modulated in Psa3 (eight induced and three repressed). Of the 75 identified HKs and HHKs, 37 contained one or more transmembrane domains (maximum = 13) and eight included a signal peptide. Various other domains were also present in these proteins (e.g., GAF, PP2C, 2CSK_N, dCache1, CHASE3, KdpD, Usp, KinB, PHY, CheW, Hypoth_Ymh and RsbRD) although the most widely represented domains were PAS/PAC (27 proteins) and HAMP (19 proteins).

**Fig 11.**
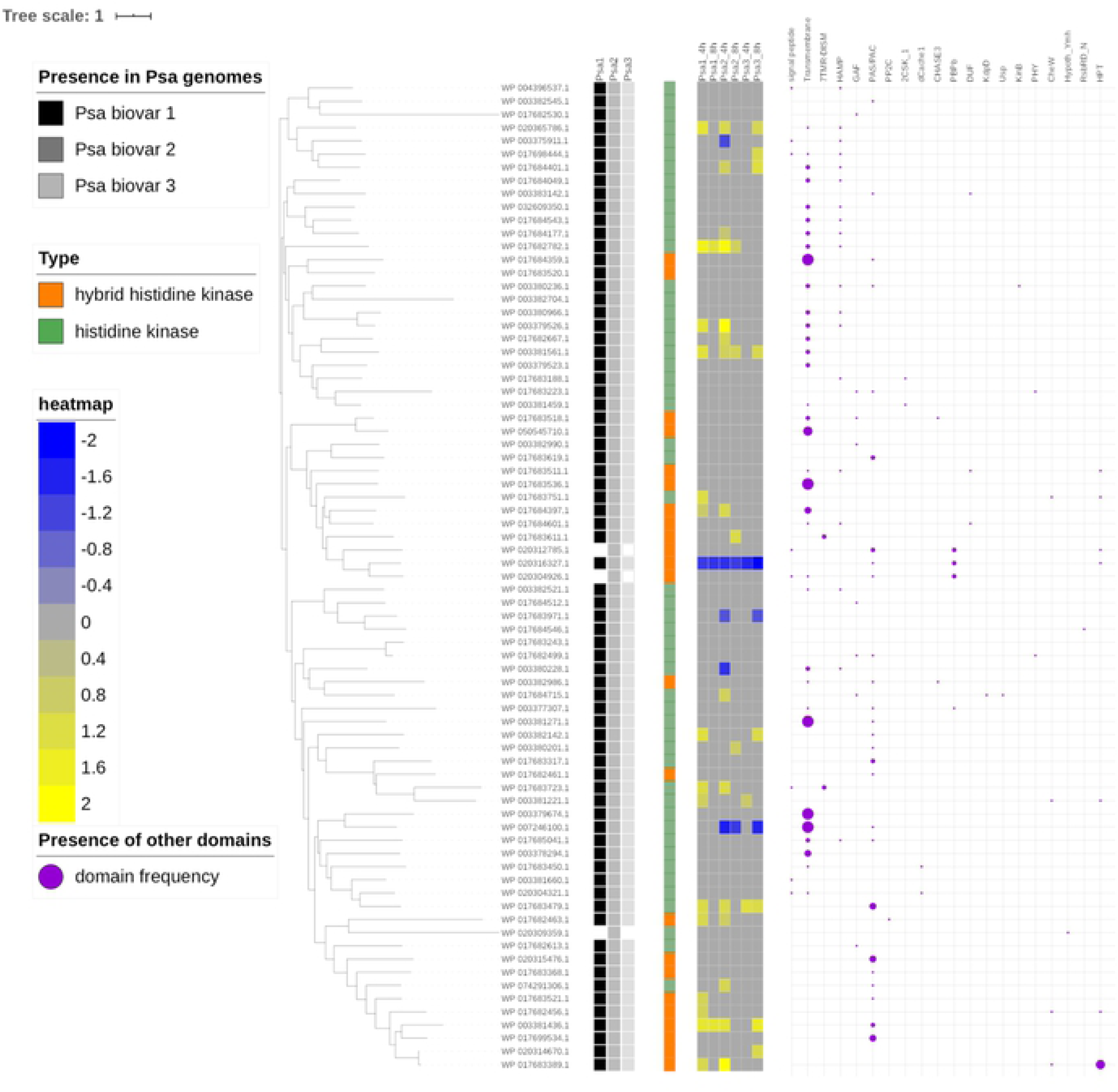
Phylogenetic analysis, expression level and main features of histidine kinases in Psa. The unrooted tree was generated using phylogeny.fr (http://www.phylogeny.fr) with the full-length protein sequences of the 75 histidine kinases identified in the Psa genomes. Branch support values were obtained from 100 bootstrap replicates. The presence of each histidine kinase (HK = green square) or hybrid histidine kinase (HHK = orange square) gene in a given genome is indicated as a colored square (black = Psa1, dark gray = Psa2, light gray = Psa3). The asterisks indicate apparent pseudogenes. The level of differential expression between minimal and rich media in the different Psa biovars is indicated by the color scale (yellow = upregulation, blue = downregulation). Psa3 is represented by the strain ICMP18884. The presence of a signal peptide, transmembrane domain or other known functional domains identified using SMART is shown on the right. The size of purple circles is proportional to the domain frequency.

Signal perception requires the presence of sensors at the onset of environmental changes, so it is tempting to speculate that basal expression levels of HK/HHK proteins (i.e., in rich medium) may also account for the specific response of different biovars to minimal medium. The hierarchical clustering of HK/HHK expression profiles using normalized fluorescence intensities, representing cells grown in rich medium, yielded one cluster containing Psa1 and Psa2 at both time points, and a second cluster containing Psa3 at both time points. These results suggest that HK/HHK expression profiles in Psa1 and Psa2 are similar in rich medium (at least for the first 8 h). In contrast, Psa1 and Psa3 clustered together and were separated from Psa2 in minimal medium, further supporting the unique behavior of Psa2 under apoplast-like conditions.

We identified eight major gene clusters with specific expression profiles in the different biovars (**Fig 12A**). Cluster 1 included HK/HHK genes that were constitutively expressed at high levels in Psa3 and at low levels in Psa2, and induced by minimal medium in Psa1 (**Fig 12B**). Cluster 7 included HK/HHK genes that showed constitutively higher expression levels in Psa3 in rich or minimal medium compared to the other two biovars (**Fig 12C**). The corresponding proteins, whose expression was not modulated by switching from rich to minimal medium, are interesting candidates for external signal perception in the most virulent biovar, potentially accounting for the rapid induction of virulence given their constitutively high expression level. Cluster 7 also included one gene that was expressed at particularly high levels in Psa3 under all conditions, but it was annotated as a pseudogene due to the presence of a small deletion producing a slightly truncated product compared to its counterpart in Psa2 (WP_020312785.1). The presence of transcripts encoded by this pseudogene was confirmed by RT-PCR (**Fig S13**).

**Fig 12.**
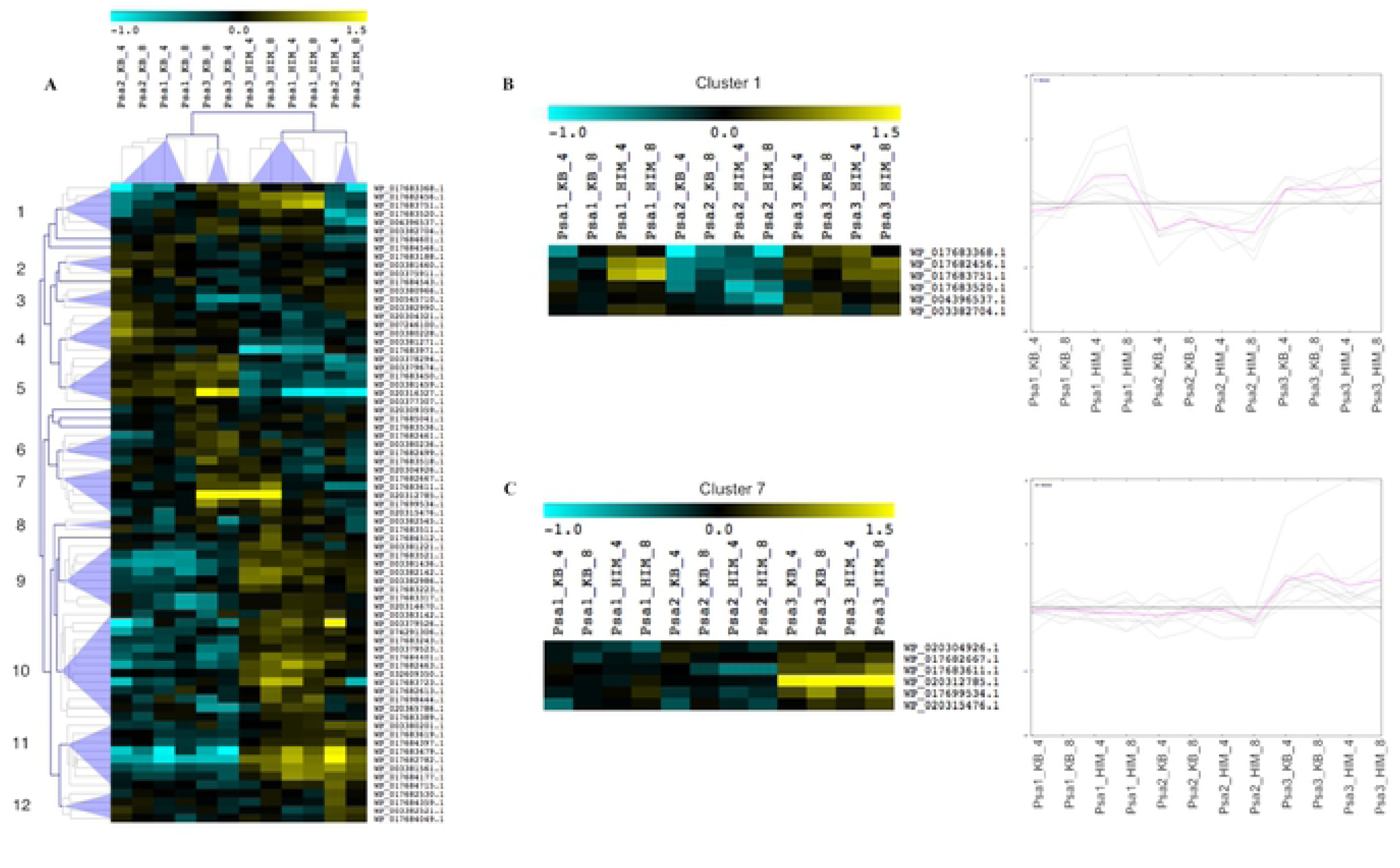
Hierarchical clustering of the expression profiles of histidine kinases in Psa. (A) Gene and sample trees obtained from the hierarchical clustering of the absolute expression level of all members of the histidine kinase (HK) family identified in this study, including hybrid histidine kinases (HHK). Normalized expression values based on microarray data were used for hierarchical clustering based on Pearson’s distance metric. The color scale represents higher (yellow) or lower (blue) expression levels with respect to the median transcript abundance of each gene across all samples. Psa3 is represented by the strain ICMP18884. Gene and sample clustering was performed with MeV using a distance threshold adjustment of 1.5. Blue triangles indicate different clusters of HK genes showing unique expression trends among Psa biovars depending on the time and growth conditions. (B) Detailed description of cluster 1 and the corresponding graphical representations of expression trends among Psa biovars. Cluster 1 = genes expressed at higher levels in Psa3 than the other strains in both media, and induced in Psa1 in minimal medium. (C) Detailed description of cluster 7 and the corresponding graphical representations of expression trends among Psa biovars. Cluster 7 = genes expressed at higher levels in Psa3 than other strains in both media and not modulated in the other strains. Samples are not clustered in (B) and (C) and the order therefore differs from the main cluster shown in (A).

The procedure described above for HK/HHK genes was repeated for the identification of RR candidates. This led to the identification of 74 common RR genes, together with one uniquely found in Psa2 and one found in Psa2 and Psa3 but not Psa1 (**Fig 13**). Any sequences containing HK domains were excluded because they were already counted among the HHK genes discussed above. Only one of the RR candidates contained a transmembrane domain, but various other domains were identified (e.g., PAS/PAC, HPT, CheW, AAA ATPase, HTH_8 (Fis-type), GGDEF, EAL, HD-GYP, CheB_methylest, CHASE3, LyTR, PP2C_SIG, TPR and ANTAR) although the most widely represented domains were Trans_reg_C (20 proteins) and HTH_LuxR (12 proteins). This was consistent with the function of RRs, which often act as direct transcriptional regulators. As already observed for the HK/HHK genes, the RR genes were mainly upregulated in response to minimal medium, with only three genes repressed (one in both Psa1 and Psa3, one in Psa2 alone, and one in Psa3 alone). Although Psa3 is the most virulent strain, only 12 RR genes were upregulated in response to minimal medium (mainly at 8 h), compared to 20 in Psa1 and 21 in Psa2 (mainly at 4 h). As discussed above for HK/HHK genes, the RR genes involved in rapid external signal transduction may be already expressed in Psa3 before the switch to minimal medium. The analysis of normalized fluorescence values revealed RR genes that were not modulated in Psa3 but were expressed at higher levels compared to the other biovars (**Fig 14**). The RR genes formed 10 distinct clusters, with cluster 1 comprising those expressed at higher levels in Psa3 than the other biovars even in rich medium (**Fig 14A**). Interestingly, some of these genes were slightly upregulated in Psa1 grown in minimal medium, but expression levels remained low in Psa2 under all conditions (**Fig 14B**). Within this cluster, the gene encoding protein WP_017682831.1 was constitutively expressed at a high level in Psa3. This gene is not present in the Psa1 genome, and is found close to one of the HHK pseudogenes in Psa3, in a chromosomal region that has been newly acquired by the most recent biovars (Psa2 and Psa3).

**Fig 13.**
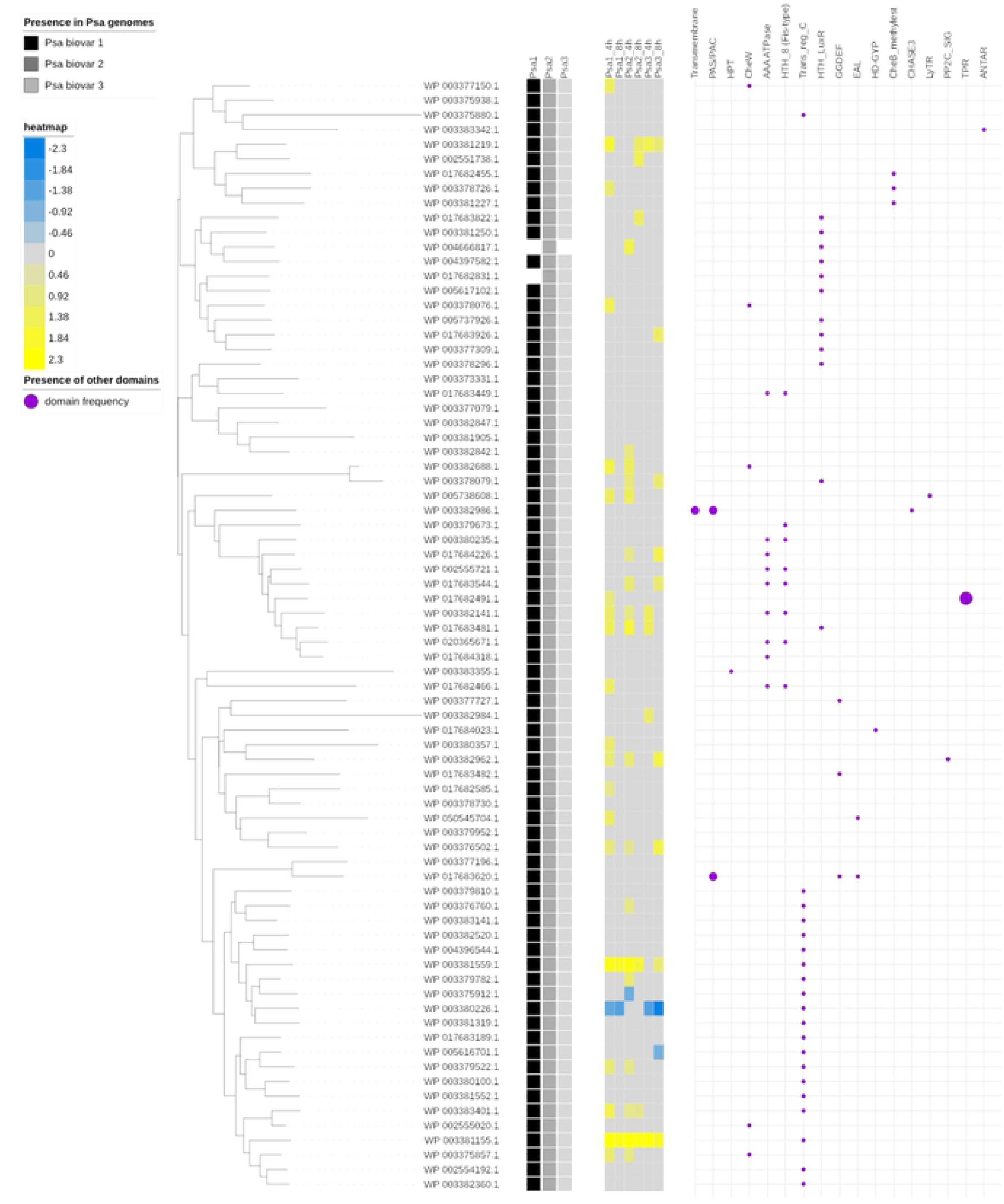
Phylogenetic analysis, expression level and main features of response regulators in Psa. The unrooted tree was generated using phylogeny.fr (http://www.phylogeny.fr) with the full-length protein sequences of the 76 response regulators identified in the Psa genomes. Branch support values were obtained from 100 bootstrap replicates. The presence of response regulator gene in a given genome is indicated as a colored square (black = Psa1, dark gray = Psa2, light gray = Psa3). The level of differential expression between minimal and rich media in the different Psa biovars is indicated by the color scale (yellow = upregulation, blue = downregulation). Psa3 is represented by the strain ICMP18884. The presence of a signal peptide, transmembrane domain or other known functional domains identified using SMART is shown on the right. The size of purple circles is proportional to the domain frequency.

**Fig 14.**
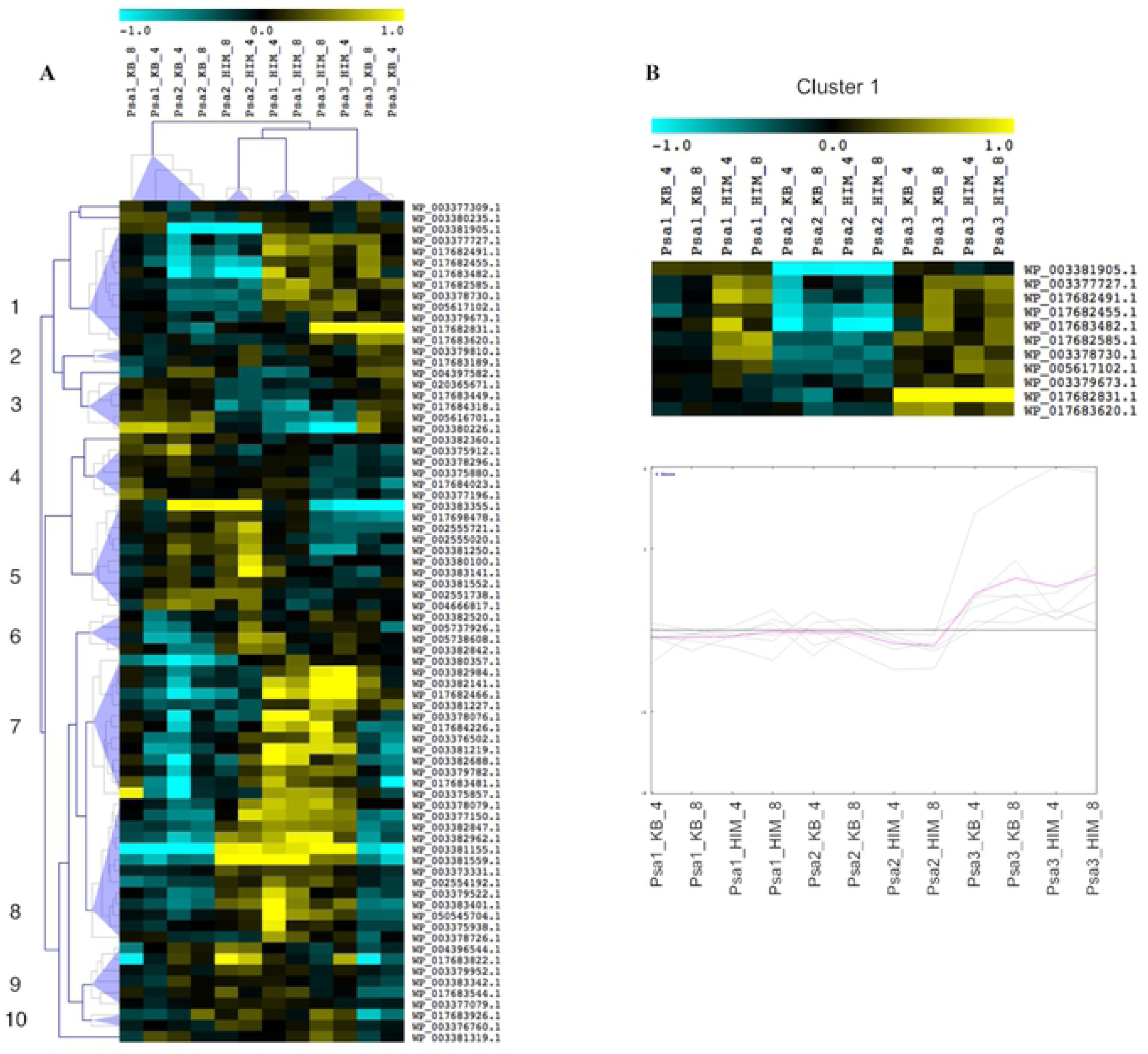
Hierarchical clustering of the expression profiles of response regulators in Psa. (A) Gene and sample trees obtained from the hierarchical clustering of the absolute expression level of all members of the response regulator family identified in this study. Normalized expression values based on microarray data were used for hierarchical clustering based on Pearson’s distance metric. The color scale represents higher (yellow) or lower (blue) expression levels with respect to the median transcript abundance of each gene across all samples. Psa3 is represented by the strain ICMP18884. Gene and sample clustering was performed with MeV using a distance threshold adjustment of 1.5. Blue triangles indicate different clusters of RR members showing unique expression trends among Psa biovars depending on the time and growth conditions. (B) Detailed description of cluster 1 and the corresponding graphical representations of expression trends among Psa biovars. Cluster 1 = genes expressed at higher levels in Psa3 than other strains in both media and not modulated in the other strains. Samples are not clustered in (B) and the order therefore differs from the main cluster shown in (A).

HKs are also regulated at the post-translational level by the second messenger c-di-GMP, which plays a key role in bacterial signal transduction (13). This molecule is produced by diguanylate cyclases, which possess a canonical diguanylate cyclase domain (GGDEF), and is degraded by phosphodiesterases (PDEs), which may possess EAL or HD-GYP domains (30). Such proteins comprise the second-most common group of two-component system output domains, establishing a link between the two major systems in bacterial signaling. HHKs are activated by c-di-GMP, which binds to a specific pseudo-Rec domain (31). We therefore carried out a whole-genome screen for GGDEF, EAL and HD-GYP domain-containing genes and analyzed their expression as above. We identified 46 genes potentially involved in c-di-GMP metabolism, 17 encoding GGDEF proteins, four encoding EAL proteins, 17 encoding proteins with both domains, and seven encoding proteins with a HD-GYP domain (**Fig 15**). Forty of the genes were common to all strains, and the remaining six were missing in at least one biovar. Nineteen of the encoded proteins contained at least one transmembrane domain and 10 also contained a PAS/PAC domain. Other domains were present in the GGDEF/EAL proteins, including pseudo-Rec (see above), dCache1, CHASE, MASE2, GAF, MHYT and HAMP. In contrast, the HD-HYP proteins contained NTP_transf_2, GlnD_UR_UTase and ACT, PolyA_Pol and PolyA_pol_RNAbd, and tRNA_edit domains.

**Fig 15.**
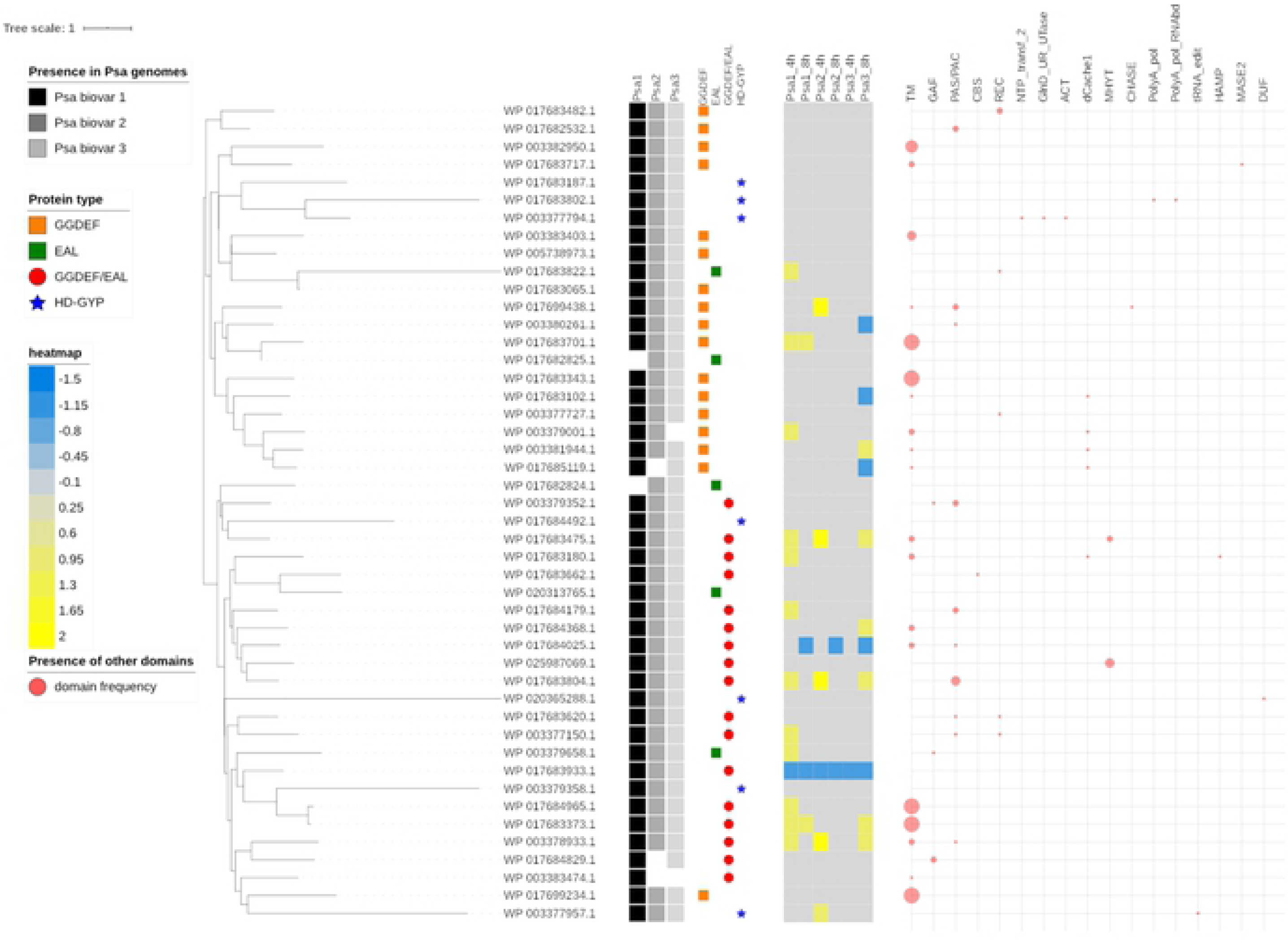
Phylogenetic analysis, expression level and main features of c-di-GMP-related enzymes in Psa. The unrooted tree was generated using phylogeny.fr (http://www.phylogeny.fr) with the full-length protein sequences of the 46 proteins containing GGDEF, EAL and/or HD-GYP domains identified in the Psa genomes. Branch support values were obtained from 100 bootstrap replicates. The presence of each GGDEF (orange square), EAL (green square), GGDEF/EAL (red circle) and HD-GYP (blue star) domain in a given genome is indicated as a colored square (black = Psa1, dark gray = Psa2, light gray = Psa3). The asterisks indicate apparent pseudogenes. The level of differential expression between minimal and rich media in the different Psa biovars is indicated by the color scale (yellow = upregulation, blue = downregulation). Psa3 is represented by the strain ICMP18884. The presence of a signal peptide, transmembrane domain or other known functional domains identified using SMART is shown on the right. The size of purple circles is proportional to the domain frequency.

Five of the genes encoding GGDEF/EAL proteins were commonly modulated in the different Psa biovars in response to minimal medium: three upregulated (WP_003378933.1, WP_017683475.1 and WP_017683804.1) and two downregulated (WP_017683933.1 and WP_017684025.1). However, most of the genes related to c-di-GMP signaling were modulated in a biovar-specific manner. Three of four GGDEF genes modulated in Psa3 were downregulated, whereas two GGDEF genes were upregulated in Psa1 and Psa2 (although different genes in each biovar). Only two EAL genes were modulated in response to minimal conditions and were uniquely upregulated in Psa1. These data point to a trend of declining c-di-GMP levels in Psa1 and Psa3 (suppression of synthesis or promotion of degradation) whereas the upregulation of genes involved in c-di-GMP synthesis tended to maintain steady levels in Psa2. Hierarchical clustering of normalized fluorescence values revealed six gene clusters with transcript profiles differing among the three biovars (**Fig 16A**). Cluster 1 contained seven genes with constitutively higher expression levels in Psa3 compared to the other biovars. Cluster 3 contained four genes with constitutively lower expression levels in Psa3 compared to the other biovars, and these genes were induced in response to minimal medium in Psa1 but not modulated by the medium in Psa2 (**Fig 16B**). The two genes from cluster 1 showing the highest expression levels in Psa3 encoded one EAL protein and one HD-GYP protein, both involved in the degradation of c-di-GMP, whereas the gene in cluster 3 most strongly repressed in Psa3 encoded a GGDEF protein involved in c-di-GMP synthesis (**Fig 16B**). The behavior of the Psa biovars was reflected at the level of sample clustering, with Psa1 and Psa2 forming one cluster in rich medium (indicating similar expression of genes involved in c-di-GMP metabolism) but resolving to different clusters following the switch to minimal medium, whereas Psa3 formed a single cluster encompassing both media (indicating the limited effect of growth conditions on the regulation of these genes). Taken together, these data indicate that Psa3 has a lower constitutive level of c-di-GMP than the other biovars, but that the high levels present in Psa1 are depleted following exposure to apoplast-like conditions whereas those in Psa2 are unaffected.

**Fig 16.**
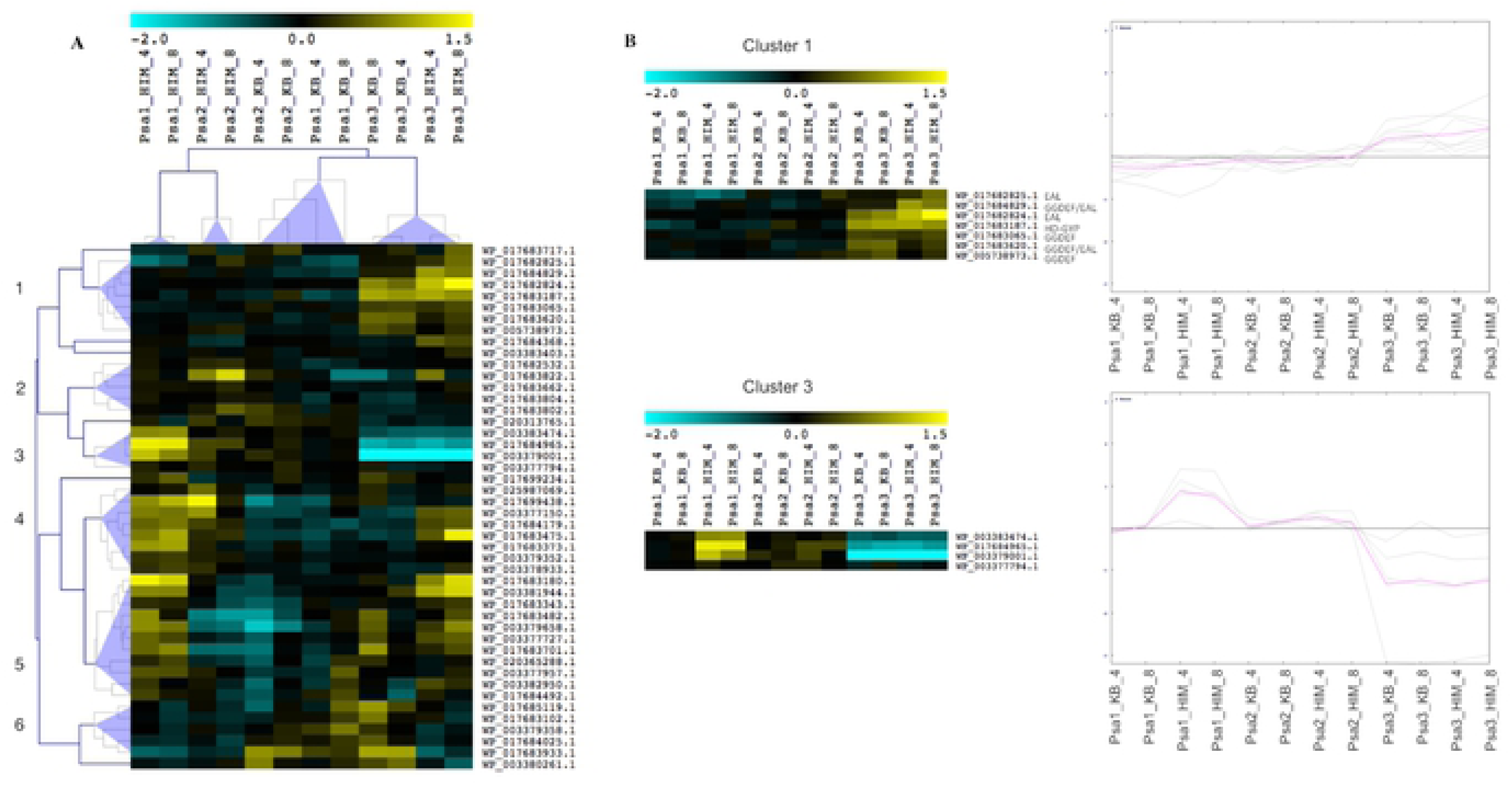
Hierarchical clustering of the expression profiles of c-di-GMP-related enzymes in Psa. (A) Gene and sample trees obtained from the hierarchical clustering of the absolute expression level of all genes containing GGDEF/EAL/HD-GYP domains identified in this study. Normalized expression values based on microarray data were used for hierarchical cluster analysis based on Pearson’s distance metric. The color scale represents higher (yellow) or lower (blue) expression levels with respect to the median transcript abundance of each gene across all samples. Psa3 is represented by the strain ICMP18884. Gene and sample clustering was performed with MeV using a distance threshold adjustment of 1.5. Blue triangles indicate different clusters of genes containing GGDEF/EAL/HD-GYP domains showing unique expression trends among Psa biovars depending on the time and growth conditions. (B) Detailed description of cluster 1 and the corresponding graphical representations of expression trend among Psa biovars. Cluster 1 = genes expressed at higher levels in Psa3 compared with other strains in both media and not modulated in other strains. (C) Detailed description of cluster 3 and the corresponding graphical representations of expression trends among Psa biovars. Cluster 3 = genes expressed at lower levels in Psa3 than other strains in both media and upregulated in Psa1. Samples are not clustered in (B) and (C) and the order therefore differs from the main cluster shown in (A).

### Inverse relationship between TTSS induction and biofilm production

Biofilm formation is positively regulated by c-di-GMP (32). Our expression data indicated that Psa2 may produce more c-di-GMP than Psa1 and Psa3, explaining the absence of TTSS induction in this biovar. We therefore tested for biofilm formation in the three strains, anticipating that Psa2 would readily form biofilms but the other biovars would not. As expected, we found that Psa1 and Psa3 were poor biofilm producers in both media, reaching maximum crystal violet absorbance values of 0.8 and 0.4 after 24 h in rich and minimal medium, respectively. Psa2 formed biofilms more readily, achieving absorbance values of 1.6 in rich medium (two-fold higher) and 1.2 in minimal medium (three-fold higher) over the same period (**Fig 17**). Differences were already visible after only 4 h, with absorbance values of 0.6 (Psa2) and 0.2 (Psa1/Psa3) in both media.

**Fig 17.**
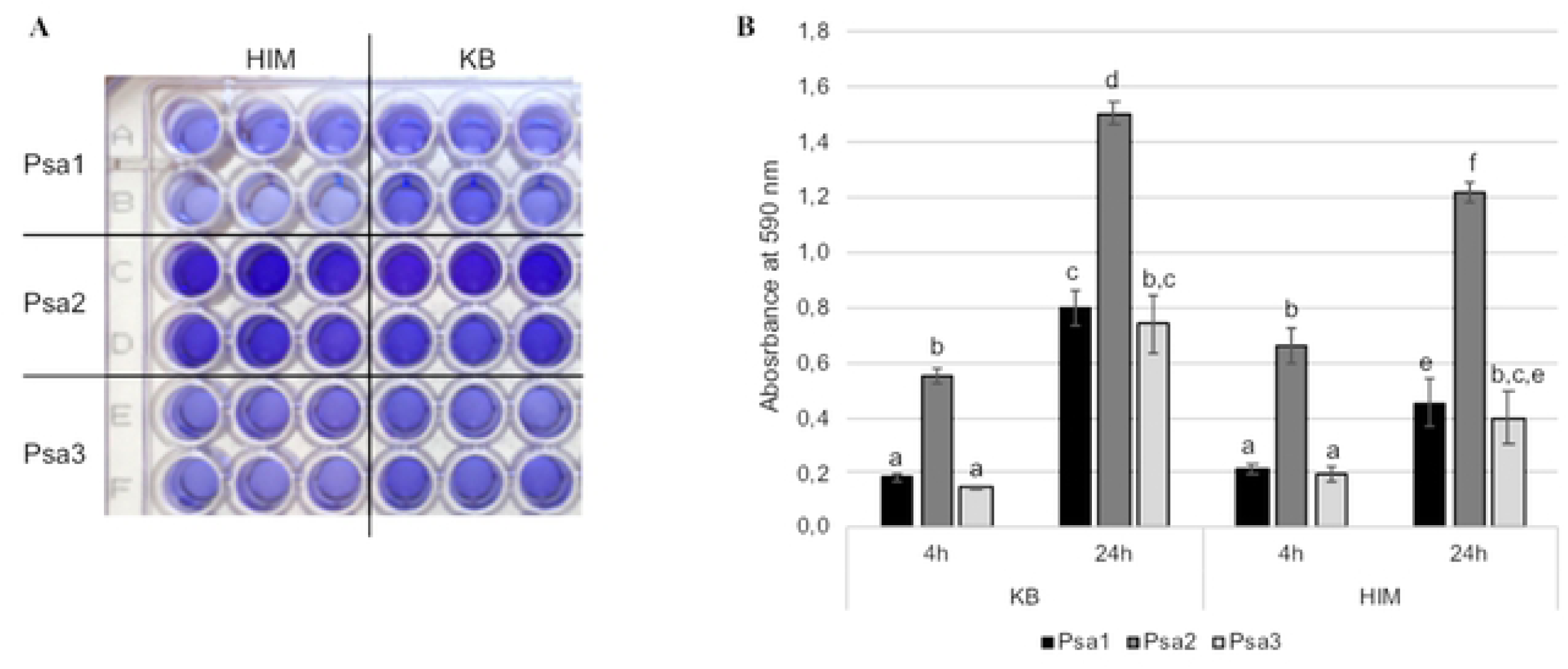
Biofilm formation in different Psa biovars. Biofilm formation was quantified in Psa1 (ICMP9617), Psa2 (KN.2) and Psa3 (CRAFRU8.43) by crystal violet staining. (A) Image of a microtiter plate following crystal violet staining of Psa1, Psa2 and Psa3 grown for 24 h in *hrp*-inducing medium (HIM) or rich medium (KB) at 28°C. For each condition, six technical replicates are shown corresponding to one of three independent biological replicates. (B) Quantification of biofilm formation in Psa cell suspensions cultured in 96-well plates for 24 h at 28°C in KB medium or HIM and stained with crystal violet. Absorbance was measured at OD_570_. Wells containing no cells were used as negative controls. Different letters indicate a statistically significant difference (p < 0.05).

## Discussion

*Pseudomonas syringae* causes frequent epidemic diseases in herbaceous and woody crops and is among the best-studied bacterial phytopathogens (33,34). Psa has recently emerged as a model organism for the analysis of bacterial pathogens affecting woody species due to the severe impact of recent disease outbreaks (35). However, it is unclear why certain Psa strains are so virulent, particularly those representing biovar Psa3 (3). We therefore investigated the aggressiveness of the main three Psa biovars using a multi-strain microarray, which remains a useful approach despite the advance of RNA-Seq technology because it provides a direct visual signature for the comparison of multiple samples. Intrinsic redundancy due to cross-hybridization was avoided by whole-genome comparisons, allowing the microarray to be used simultaneously for transcriptomic analysis and the tentative functional annotation of genes not yet represented in the available genome sequences of poorly-characterized strains. Our single multi-strain microarray also provided a common reference name for all Psa strain genes and orthologs from the control pathovar Pto.

### Psa biovars and Pto share a common response to nutrient deficiency

The microarray data revealed the common upregulation of many genes involved in transcriptional regulation as a core mechanism shared among the Psa and Pto strains. The modulated genes included those encoding sigma factors (which control the transcription of broad sets of target genes) and proteins related to phosphate starvation, including a PhoH-like protein, the transcriptional regulator PhoP, the phosphate starvation-inducible protein PsiF, and the phosphonate metabolism protein PhnM. Phosphate limitation has a pleiotropic effect on bacterial physiology, triggering the degradation of polyphosphate, the accumulation of phosphate-free membrane lipids, and the accumulation of polyhydroxyalkanoate (PHA) in regulatory granules (36,37). The Psa biovars and Pto induced two main genes related to PHA synthesis, encoding the phasins PhaI and PhaF. However, the functional category related to PHA appeared enriched only in Psa1, which induced three other PHA-related genes despite the generally low number of genes uniquely upregulated in this biovar. We also observed the strong common induction of a MaoC-like hydratase (MaoC), an (*R*)-hydratase that links β-oxidation to the PHA biosynthesis pathways (38–41). PHAs provide intracellular carbon and energy reserves under nutrient-limited conditions with excess carbon (37), so this pathway may provide a common adaptive response allowing *P. syringae* to overcome nutrient limitation within the apoplast.

The stringent response triggers the downregulation of genes encoding translational components and the simultaneous upregulation of genes involved in amino acid biosynthesis and transport (42). This reflects the combined actions of the DnaK suppressor protein DksA and the nucleotide alarmones (p)ppGpp on RNA polymerase (43). Accordingly, we observed the induction of two genes encoding DnaK suppressor proteins but we did not observe the modulation of genes related to stress-induced alarmone production (RelA, SpoT). All strains also induced an operon encoding the serine protein kinase PrkA, a protein containing a von Willebrand Factor type A domain, and a SpoVR-like protein (44) that has been implicated in host-pathogen interactions (45).

One of the most strongly induced genes in all strains (including Pto), with a log_2_FC > 5, encoded a C1 family papain-like cysteine protease, which potentially regulates the activity of virulence factors. Secreted proteases can help pathogens avoid recognition by the plant immune systems, and many TTSS-dependent effector proteins in Pto (e.g., AvrRpt2, AvrPphB, HopPtoN and HopX) are clan CA proteases, which share the same papain-like 3D structure, order of catalytic residues, and other conserved features (46–48). However, the candidate we identified is unlikely to be an effector because it was also strongly induced in Psa2, in which the TTSS is inactive. Given the importance of proteases for protein quality control systems (49), this protease may be a stress-response protein that allows Psa and Pto to adapt in the hostile host environment, or it may be active in the apoplast.

### Psa biovars also show unique responses to apoplast-like conditions

The microarray data also revealed many biovar-specific responses to nutrient restriction. In Psa1 and Pto, we observed a significant upregulation of chemotaxis-related genes. This contrasts with a previous study showing the repression of such genes in *P. syringae* pv. *syringae* B728a in the apoplast environment, probably reflecting differences in the media used in these experiments (50). Two chemotaxis pathways (*che1* and *che2*) are required for the complete fitness of Pto but primarily function during the epiphytic phase, because the corresponding mutants cannot enter the apoplast (51). However, this does not rule out the possibility that chemotaxis is also required for virulence, allowing bacteria to move within the apoplast to colonize the host more efficiently. The main mechanism associated with chemotactic control is flagellum-related motility, which was induced in Pto, Psa1 and Psa3 but not Psa2. Among the motile strains, Psa1 showed the most significant enrichment in flagellum-related genes, and the HKs WP_017683751.1 and WP_017682456.1 (present in all strains) were specifically induced only in this biovar. Both HKs are located in operons also containing genes related to flagellum assembly and chemotaxis. The specific induction of these HKs may account for Psa1 motility in response to minimal medium. In Psa2, the inability to induce genes encoding flagellum components may reflect the upregulation of sigma factor AlgU, a repressor of flagellar gene expression *in planta* (52). This may avoid host immune responses because flagellum components act as pathogen-associated molecular patterns (PAMPs).

In Psa3 (and Pto), we also observed the induction of genes related to iron transport and chelation, in particular several genes involved in pyoverdine synthesis. Iron is required for many key metabolic functions in bacteria, including the tricarboxylic acid cycle, electron transport chain, and DNA synthesis (53), and it also induces several virulence genes in Pto (54). However, iron levels in the apoplast do not limit Pto growth (55) and PvdS-regulated iron-scavenging systems are not required for Pto pathogenesis (56) suggesting that iron scavenging does not account for the aggressiveness of Psa3. The induction of *anmK*, encoding an anhydro-1,6-muramic acid kinase involved in cell wall metabolism, may contribute to the virulence of Psa3 given its similar role in Pto (57).

In Psa2, we observed the induction of genes related to branched-chain amino acid metabolism, including those required for the synthesis of core metabolic precursors such as pyruvate, acetyl-CoA and oxaloacetate. Branched-chain amino acids are important nutrients but also act as signals that induce virulence gene expression, which in some Gram-negative bacteria is regulated via the leucine-responsive regulatory protein Lrp (58,59). The Psa2 *lrp* gene was not modulated by minimal medium, suggesting that branched-chain amino acids regulate virulence via other pathways, as shown for *Xanthomonas oryzae* pv. *oryzae* (60). Leucine and valine are also precursors for the synthesis of diffusible signal factors (DSFs), which act as quorum sensing molecules. Several genes related to leucine and malonate metabolism were upregulated in Psa2, including crotonase/enoyl-CoA hydratase, malonate decarboxylase subunit gamma, the malonate transporter subunit MadL and a malonyl CoA-acyl carrier protein (ACP) transacylase. This suggests the induction of malonyl-ACP synthesis, which may feed into the fatty acid synthase (FAS) elongation cycle to generate DSF precursors (61). Accordingly, we also observed the upregulation of the FAS gene *fabG* encoding β-ketoacyl-ACP reductase specifically in Psa2. Given that quorum sensing signals have not yet been identified in Psa, our data provide tantalizing evidence that DSFs may fulfil this role in Psa2 (4,62,63).

The Psa2 genome is unique among the Psa biovars because it carries a complete metabolic pathway for the synthesis of coronatine (12) and these toxins have been synthesized in optimized medium at 18°C (64). Interestingly, we did not observe the modulation of coronatine-related gene expression under our apoplast-like conditions in Psa2, although these genes were modulated in Pto. However, the *corR* gene, encoding a positive regulator of coronatine biosynthesis, was strongly induced by minimal medium in Psa2 but not in Pto. In a previous study, the Pto *corR* gene was slightly induced in both Hoitink-Sinden medium supplemented with sucrose and *hrp*-derepressing medium (65), suggesting that induction was below the limit of detection in our experiments. Conversely, the expression of *corS*, encoding the cognate HK, was significantly lower in Psa2 than Pto. Both elements of the two-component CorRS system are required for coronatine biosynthesis (66), so the low expression of *corS* in Psa2 may be insufficient to switch on coronatine metabolism. The differential expression of *corR* and *corS* may reflect their relative positions in the *corRS* operon, as described for certain structured operons in other bacteria (67)(68). Furthermore, the operon may be regulated by external factors, as reported for the temperature-dependent regulation of coronatine synthesis in the soybean pathogen *P. syringae* pv. *glycinea* PG4180 (Psg), whereas coronatine production in Pto is temperature-independent (69,70). Given the 99% sequence identity between the CorS proteins in Psa2 and Psg, it is tempting to hypothesize that coronatine synthesis in Psa2 is also temperature-dependent, and that our experimental conditions (28°C) do not favor this metabolic pathway.

### Genomic analysis reveals new features of the TTSS-related pathogenicity island

The TTSS plays a key role in Pseudomonas–host interactions, and an activating signal is required to induce the secretion of virulence effectors (16) that suppress the host immune system and allow the bacteria to establish a niche in the apoplast (71). Although the signals are poorly understood, there is compelling evidence that TTSS activation occurs upon contact with target cells (13,14) and that the *hrp*/*hrc* gene cluster is controlled by multiple physiological and environmental factors that are replicated by minimal medium (15). We therefore anticipated the significant induction of *hrp*/*hrc* genes in Pto, as reported previously (15,72,73). We also observed the same phenomenon in Psa3, whereas the *hrp*/*hrc* genes in Psa1 were induced weakly and those in Psa2 were not induced at all. These data suggest that different biovars of the same pathovar respond in different ways to the apoplast-like environment and may therefore rely on different pathogenicity strategies.

A comparison of the complete 23,721-bp *hrp*/*hrc* cluster revealed high cross-strain conservation, except for a slight divergence within the *hrcU* operon in Psa2 compared to Psa1 and Psa3, but this did not appear to affect the promoter regions or gene products. The *hrp*/*hrc* cluster showed 99.94% identity between Psa1 and Psa3 and 99.46% identity between Psa2 and Psa1/Psa3, not including the gap in Psa2 corresponding to the hypervariable intergenic region. Focusing on the *hrcU* operon, the identity between Psa2 and the other biovars decreased to 97.6%. Overall, the *hrp*/*hrc* cluster showed a canonical structure delimitated by *hrpR* and *hrpK* in all biovars, and a similar structure in Pto (8). However, the detection of a polycistronic transcript featuring *hrpO* and *hrpP* suggests that the current organizational model of the *hrpU* and *hrpJ* operons should be revised, and the presence of the *hrp* box within the coding sequence of *hrpO*, the first gene in the operon, suggests a novel mode of operon regulation that may be more flexible (74). Our data suggest that the *hrp* box within the *hrpO* coding sequence may not necessarily regulate *hrpP* but could facilitate the regulation of *hrpO* and downstream genes by the sigma factor HrpL.

Another key feature of TTSS-containing pathogenicity islands is the presence of an exchangeable effector locus (EEL) downstream of the *hrp* cluster (8). Our Psa2 and Psa3 strains featured the canonical EEL structure (including tRNA^Leu^, *queA* and *tgt* sequences) although two of the three Psa2 genomes (ICMP 19071 and ICMP 19072) contained a phage insertion within the EEL, supporting the observation that tRNA genes are often found at phage integration sites (75). Multiple rearrangements have been reported in Psa1 and Psa3 biovars (76), and accordingly we found that the tRNA^Leu^, *queA* and *tgt* sequences were dispersed to other chromosomal locations in Psa1 (ICMP9617 and ICMP9853). Such rearrangements may be selectively neutral, but the genome structure may affect transcriptional regulation (76). Given that the EEL pays a role in bacterial fitness (8), it is tempting to speculate that the lower virulence of Psa1 compared to Psa3 may in part reflect the loss of the canonical EEL structure.

### The modulation of positive TTSS regulators does not correlate with the induction of *hrp*/*hrc* genes in different Psa biovars

The three Psa biovars contained similar pathogenicity islands yet the *hrp*/*hrc* gene cluster in Psa2 was not induced. This primarily depends on HrpL, a master regulator of bacterial TTSS genes (77). The *corRS* operon contains a HrpL-dependent promoter, and accordingly coronatine production is abolished in Pto *hrpL* mutants (65,78). The absence of *hrpL* expression in Psa2 may therefore contribute to the lack of coronatine-related gene expression described above. However, CorR is a positive regulator of the *hrp*/*hrc* regulon in *P. syringae* (65) and a Pto *corR* mutant showed a reduction and delay in the expression of *hrpL* and an impaired hypersensitive response on *Nicotiana benthamiana*. One hypothesis to explain the absence of CorRS signaling in Psa2 is the suboptimal temperature, which might prevent the induction of *hrpL*, but the *hrpA1* promoter was not activated in Psa2 even at 24°C or 18°C, thus refuting this hypothesis (**Fig S14**).

Intriguingly, many upstream regulators of the TTSS were strongly induced in Psa2 after 4 h in minimal medium (log_2_FC > 1.5, adj. p-value < 0.05) but modulation in Psa1 was negligible and modulation in Psa3 was only observed after 8 h. Our analysis included AefR (79), AlgU, which regulates *hrpL* and *hrpRS* directly (52), and CsvR, a HK that cooperates with CsvS to induce *hrpRS* (80). Moreover, the *hrpA1* gene is required for full expression of both secreted proteins (HrpW) and components of the Hrp secretion machinery (HrcC and HrcJ) in minimal medium and *in planta* (81). Not only was the *hrpA1* gene not induced in Psa2, but the transcript was up to eight-fold less abundant compared to the other biovars in minimal medium and up to three-fold less abundant in rich medium, and was expressed at significantly lower levels than all other *hrp* genes located in the same operon except *hrpZ1*. These data suggest that *hrp*/*hrc* cluster genes may be repressed in Psa2, due to the low expression of *hrpA1*. Surprisingly, *hrpT* was upregulated in Psa2 even though it should be dependent on HrpL binding to the *hrp* box controlling the *hrpC* operon, suggesting an alternative and independent form of transcriptional regulation. However, HrpT inhibits the expression of *hrpL* indirectly (82,83), so the induction of *hrpT* may also explain the repression of the TTSS in Psa2.

The induction of TTSS regulators in Psa3 did not fit with the strong induction of *hrp*/*hrc* cluster genes already observed at 4 h in this biovar. However, the strong expression of these genes in rich medium indicated they were already upregulated during overnight growth of the inoculum. Indeed, after >20 h the rich medium is depleted of nutrients and behaves similarly to minimal medium, which may trigger TTSS-related responses to nutrient deficiency in Psa3. This is supported by the declining TTSS-related transcript levels in rich medium after 8 h. On the other hand, the massive induction of these genes in minimal medium agrees with the perception of nutrient-independent signals in the apoplast, such as acidic pH, triggering a more profound TTSS response. The ability of Psa3 to integrate multiple external signals may contribute to its virulence, enabling cells to rapidly deploy their resources for an attack on the host plant. The upregulation of *anmK* specifically in Psa3 was also notable because a nonsense mutation in *anmK* was recently shown to impair the TTSS in a Pto *gacA* mutant background (57). Moreover, the alignment of *hrpL* promoters from the different biovars revealed a single mutation in Psa3 that created a putative supplementary binding site for the transcription factor LysR (**Fig S15**). It is unclear whether this influences *hrpL* expression and TTSS activation in Psa3, but the presence of such a mutation possibly influencing the expression of a master regulator provides new insight into the TTSS-related virulence of Psa3.

### The role of two-component systems and c-di-GMP in biovar-dependent responses

Pathogenic bacteria use sophisticated two-component signaling systems in order to adapt to environmental conditions within the host and to trigger virulence effectors (84,85). We investigated the potential role of two-component systems in the virulence of Psa3, but found no evidence of HK/HHK or RR genes that were uniquely present or specifically upregulated in this biovar. However, the comparison of transcript levels revealed some HK/HHK and RR genes that were constitutively expressed at higher levels in Psa3 (i.e., in both rich and minimal medium) but not modulated in Psa1 or Psa2, or that were induced in Psa1 but not Psa2 when switched to minimal medium. In particular, two HHK genes that are missing from Psa1 and annotated as pseudogenes in Psa3 were nevertheless strongly expressed in Psa3. One of them (WP_020312785.1) featured a 5-bp deletion in Psa3 leading to a frameshift and premature stop codon compared to its ortholog in Psa2 (**Fig S16A-B**), but the promoters were conserved in both biovars. The promoter sequence conservation and high level of expression indicates the products are likely to be functional. Apparent pseudogenes have previously been shown to produce noncoding RNAs with a role in gene expression (86) or even active products corresponding to truncated proteins (87,88). The corresponding gene in Psa2 encodes an HHK similar to the *E. coli* EvgS protein, which perceives mild acidic conditions (pH 5.5) via a cytoplasmic linker (89). The translation of the Psa3 pseudogene *in silico* indicated two putative truncated proteins, the first corresponding to the periplasmic region (including the PBDb and PAS domains) and the second also containing the cytoplasmic region including three key catalytic domains (HK, HPT and REC), similar to the pH-sensing cytoplasmic portion of EvgS (**Fig S16C-D**). In *E. coli*, EvgS activates EvgA, a RR containing a helix-turn-helix LuxR DNA-binding motif (90). Accordingly, we found that a similar RR gene (WP_017682831.1) was also strongly expressed in Psa3 and was located close to the HHK-like pseudogene WP_020312785.1 discussed above, within a genomic segment absent from Psa1. Our data therefore suggest that a truncated HHK derived from this apparent pseudogene interacts with the RR to transduce a pH-dependent signal for TTSS activation in the apoplast. This agrees with evidence that pseudogenes can be called upon as a reservoir of genetic information to deal with environmental stress or conditions that promote mutation (91,92).

The transduction of signals is facilitated by second messengers such as c-di-GMP and (p)ppGpp (93,94). The synthesis of c-di-GMP requires a diguanylate cyclase containing a GGDEF domain whereas excess c-di-GMP is degraded by PDEs containing an EAL or HD-GYP domain. Pto produces one diguanylate cyclase and one PDE that modulate c-di-GMP levels to control virulence (95,96). Genetic manipulation of the c-di-GMP content has shown that the accumulation of this signal promotes biofilm production while inhibiting motility and the TTSS (32). We found 46 proteins with GGDEF and/or EAL/HD-GYP domains potentially involved in c-di-GMP metabolism and signaling. Interestingly, GGDEF-containing proteins were mostly repressed in Psa3 but not the other biovars, whereas some putative PDEs were constitutively expressed at higher levels in Psa3 than the other biovars. Two PDE genes (WP_017682825 and WP_017682824) were located just downstream of the HHK pseudogene and cognate RR gene discussed above, suggesting this region of the genome, missing in Psa1, is a key determinant of pathogenicity. In proteins with both GGDEF and EAL domains, one of the domains is usually inactive and fulfils a regulatory function, or a third regulatory domain is present to disjoin the activity of the GGDEF and EAL domains (97,98). We identified two hybrid GGDEF/EAL proteins that were strongly expressed in Psa3. One of them (WP_017684829) also included a GAF domain, suggesting that other small molecules such as c-di-GMP or cAMP can bind to the GAF domain and modulate c-di-GMP synthesis and/or degradation (99). The other (WP_017683620) also contained PAS/PAC and REC domains, suggesting it may function as a RR, thus linking the activation of a RR directly to the modulation of c-di-GMP by promoting or repressing diguanylate cyclase or PDE activity (99). These data suggest that Psa3 tends to reduce c-di-GMP levels by suppressing its synthesis and/or accelerating its degradation. Given the increased virulence of *Pseudomonas* mutants overexpressing the PDE BifA (96), the aggressiveness of Psa3 may also involve the *hrp*/*hrc*-dependent depletion of c-di-GMP leading to the induction of TTSS-dependent signaling (25,32).

In Psa2, the higher c-di-GMP content may account for the inactive TTSS and overall lower virulence. The level of c-di-GMP is regulated by the chemoreceptor PscA, and Pto *pscA* mutants accumulate this second messenger and favor biofilm formation over swarming motility (100). Although *pscA* was upregulated in Psa2 in minimal medium at 4 h, it was still expressed at higher levels in Psa3 in both media at both time points (log_2_FC > 2; adj. p-value < 0.05). This low level of *pscA* expression may contribute to the higher c-di-GMP levels in Psa2, which in turn would suppress motility-related genes such as *flhA, fliE* and *fliN* (32). Indeed, we found that these genes were expressed at lower levels in Psa2 than Psa3 (log_2_FC > 1.5, adj. p-value<0.05). The higher c-di-GMP level in Psa2 would also explain the greater propensity of this biovar for biofilm formation in both media compared to the other biovars.

Taken together, our data suggest that c-di-GMP is key marker of aggressiveness in different Psa biovars, and acts as a switch between sessile behavior (biofilm formation) and virulence (TTSS activation). Psa3 appears to maintain lower c-di-GMP levels by the constitutive expression of PDE genes and the suppression of diguanylate cyclases, whereas the expression of diguanylate cyclases in Psa2 leads to the repression of TTSS responses by c-di-GMP. In line with the intermediate phenotype we observed, Psa1 suppresses c-di-GMP-degrading enzymes containing an EAL domain when transferred to minimal medium, thus requiring more time than Psa3 to deplete any reserves of this second messenger, leading to the weaker induction of TTSS responses compared to Psa3.

## Conclusion

Our multi-strain transcriptome profiling strategy allowed us to identify subtle differences between closely-related bacterial strains under different conditions and to analyze genes that have yet to be annotated in particular strains, such as *corR* and *corS* in Psa2. More importantly, we were able to confirm the expression of pseudogenes in Psa3 that may have been excluded from analysis based solely on single-strain gene catalogs. We focused on TTSS effectors which play a key role in the fate of bacterial interactions in susceptible and resistant plants (101,102). Comparative genomics has previously identified strain-specific effectors that were assumed to confer virulence, such as those present in Psa3 but not Psa1 (11,12). However, this study highlights the importance of transcriptional analysis to confirm whether such effectors, and the corresponding TTSS components, are expressed and correlated with virulence traits. Differences in the timing of TTSS induction and secondary virulence mechanisms may provide an advantage to certain strains, increasing their aggressiveness. Plants can block the translocation of TTSS effectors although the mechanism is unclear, so the ability of bacteria to respond quickly to the apoplast environment would confer an advantage allowing them to overcome these defenses (103).

The identification of factors that determine virulence, such as specific two-component systems and corresponding signaling pathways and regulatory networks, is necessary for the development of strategies to reduce the virulence of plant pathogens. Our work shows how *P. syringae* strains belonging to the same pathovar, infecting the same host with different degrees of aggressiveness, may display profoundly different responses to the apoplast, thus accounting for biovar diversity in terms of virulence. Psa is not only a devastating kiwifruit pathogen, but also a very important model to study the regulation of bacterial TTSS activity. In particular, we present evidence that c-di-GMP may be a key factor controlling the aggressiveness of Psa biovars and further investigations should focus on the direct targets of c-di-GMP that control the biofilm/TTSS switch.

## Materials and methods

### Microarray design and fabrication

Whole-genome sequences representing four Psa strains and Pto, as well as three ICE sequences, were included in the design of an Agilent custom high-density microarray chip (Agilent Design ID: 078853; GEO accession: GPL27505). At the time, complete genome sequences were available for Psa3 ICMP18884 and Pto DC3000, and partial sequences were available for the others. The sequences and gene annotation data were collected in July 2015 from the NCBI GenBank and RefSeq databases (https://www.ncbi.nlm.nih.gov/) and the University of Udine (CRAFRU8.43 annotations from Prof. Giuseppe Firrao) (**S1 Appendix** and **S2 Appendix**). If multiple genome annotations were available for a given strain, a unique reference transcriptome was created by merging the data to avoid artificial redundancy at the strain level, giving the following priority for each transcript: RefSeq > GenBank > University of Udine (104). Non-redundant sequences annotated for each strain were then grouped by collapsing identical sequences into one representative to create a multi-strain, non-redundant collection of protein coding sequences (CDSs) covering all annotated transcripts for all strains (N=20,561 total CDSs) (**S3 Appendix**). The Agilent eArray web tool (https://earray.chem.agilent.com/earray/) was used to design probes based on genomic sequences representing each gene family using the following parameters: Method = Tm Matching Methodology; Probe length = 60 bp; Number of probes per target = 3; transcriptome details = “use target file as transcriptome”; Probe design = “design without 3′ bias”. Gene sequences were annotated using the Blast2GO suite v4.1.9 (105) after mapping against the NCBI non-redundant database (https://www.ncbi.nlm.nih.gov/refseq/about/nonredundantproteins/) with BLASTX v2.6.0 (22). The eArray design application yielded 18,598 high-performance probes (lowest possible target ambiguity) interrogating 20,554 protein sequences, with 14,457 unambiguous probes matching a unique target (**S4 Appendix**). The design procedure for the multi-strain microarray is summarized in **Fig S1**.

### Bacterial strains and growth media

Single colonies of the Psa3 strains CRA-FRU 8.43 and ICMP18884/V-13, the Psa1 strain J35, the Psa2 strain KN.2, and the Pto strain DC3000 grown on rich solid medium (KB agar) were inoculated into KB medium and incubated overnight at 28°C, shaking at 200 rpm. When cells reached the late log phase, they were collected by centrifugation (5000 × g, 10 min, room temperature), washed three times in liquid KB medium (106) or HIM (15), then resuspended at a final density of 2 × 10^8^ cfu/mL in HIM or 2 × 10^7^ cfu/mL in KB and incubated for 4 or 8 h at 28°C, shaking at 200 rpm. The cells were harvested by centrifugation as above and the pellets (∼2.4 × 10^9^ cells) were stored at −20°C. The procedure was carried out twice to obtain three independent biological replicates of each sample for microarray analysis.

### RNA extraction and microarray chip hybridization

RNA was extracted from each sample using the Spectrum Plant Total RNA Kit (Sigma-Aldrich) and quantified using a Nanodrop spectrophotometer (Thermo Fischer Scientific). RNA quality was evaluated using an Agilent RNA 6000 Nano Kit Bioanalyser. RNA was processed and labeled for microarray analysis using the One-Color Microarray-Based Gene Expression Analysis Low Input Quick Amp WT Labeling kit (Agilent Technologies), according to the manufacturer’s instructions.

### Microarray data analysis

The fluorescence intensity for each probe was measured using an Agilent G4900DA SureScan Microarray Scanner System with Agilent Scan Control software, and data were extrapolated using Agilent Feature Extraction software. Fluorescence intensities were calculated by robust multi-array averaging, including adjustment for the background intensity, log_2_ transformation, and quantile normalization. Normalized data were used to identify differentially expressed genes (DEGs) with threshold values of P < 0.05 and log_2_FC values > |1|. DEGs were compared across different strains and/or conditions using the online software Calculate and Draw Custom Venn Diagrams (http://bioinformatics.psb.ugent.be/webtools/Venn/). Annotation of gene sequences Gene Ontology (GO) terms (**S5 Appendix**) and GO functional enrichment analysis was carried out using Blast2GO v4.1.9 (FDR < 0.01).

### Gene sequence clustering for inter-strain comparisons

A non-redundant dataset of amino acid sequences (N=17,047; **S6 Appendix**) representing all proteins annotated from RefSeq or Genbank collected in this study (**Fig S1**) was created by taking each identical sequence only once. To allow inter-strain comparisons of protein sets, we have grouped the different proteins annotated for each strain of interest from different sources (**Fig S1**) based on sequence similarity and then analyzed the presence or absence of the given protein cluster across Pseudomonas strains.

Grouping of proteins was performed in two steps based on results of all-against-all alignments of amino acid sequences performed with the BLASTP software version 2.7.1 (22).

In the first grouping step, any pair of protein sequences scoring as reciprocal best hits (RBHs) in the BLASTP output was grouped in the same cluster (**S7 Appendix**). To assess the quality of this procedure, we compared clusters of proteins called in our analysis against a set of non-redundant RefSeq protein records annotated for *P. syringae* (marked by protein identifier: WP_XXXXXXX.1, where XXXXXXX is a unique numeric code) and found that all such instances annotated in multiple strains were correctly assigned to the same cluster of proteins (**S7 Appendix**).

Next, we further grouped clusters of protein sequences marked by high sequence similarity among cluster members (ie., E < 0.001; at least 94% identity over at least 90% of the aligned sequences in the BLASTP output) in super-clusters (**S7 Appendix**).

The above procedure assigned the full and redundant collection of protein sequences (N=56,042 FASTA sequences; **S2 Appendix**) collected for different *P. syringae* strains from different annotation sources (**Fig S1**) to a total of 10,839 protein groups. These protein groups were mapped to the 18,598 probes of the *P. syringae* multi-strain microarray to allow comparative transcriptomic investigation (**S7 Appendix**).

### Identification of HK/HHK and RR sequences, and genes related to c-di-GMP metabolism

HK/HHK and RR genes were retrieved using MiST v3.0 (107). Genes containing GGDEF or EAL domains were retrieved by using BLASTP to search annotated proteins in the Psa genomes with representative domain sequences from the RCSB Protein Data Bank (http://www.rcsb.org/): protein O35014 for the EAL domain, and protein B8GZM2 for the GGDEF domain. Genes containing an HD-GYP domain were similarly identified by screening with domain ID COG2206 from the CDD domain databank (http://structure.ncbi.nlm.nih.gov/Structure/cdd/cdd.shtml). Proteins that were not common to all three Psa biovars were used as BLASTP queries to search the proteins annotated in other Psa genomes to confirm biovar specificity, including biovars not included on the microarray (ICMP9853 for Psa1, ICMP19071 and ICMP19072 for Psa2, and CRAFRU8.43 for Psa3). A similar BLASTP search against the Psa genomes was carried out for all HK/HHK and RR genes previously identified in Pto (29). Finally, for proteins still absent from one or more Psa strains, the probes matching the corresponding genes were used as BLASTN queries against the complete Psa genomes to account for potential errors in genome annotation and the presence of pseudogenes, thus verifying the presence or absence of the corresponding genes at the nucleotide level.

### Phylogenetic analysis

For each protein family, an unrooted phylogenetic tree was built using the phylogeny.fr (http://www.phylogeny.fr/) “a la carte” pipeline, as previously described (108) with slight modifications. Phylogenetic trees were built using protein sequences and “alignment curation” was avoided in the pipeline due to the high variability among protein sequences. Results were downloaded in Newick format for tree representation using the Interactive Tree of Life (iTOL) v3.2.4 (http://itol.embl.de/). SMART (109,110) was used to identify domain architecture and protein motifs with default parameters. The dataset was uploaded for the visualization of protein classification (HK, HHK, GGDEF, EAL, and HD-GYP) and the presence of other domains using iTOL.

### Hierarchical clustering

Normalized fluorescence intensities from microarray experiments were imported as data matrices into MeV (111). The data were adjusted as median center genes/rows and clustered using the hierarchical clustering module. Gene and sample trees were clustered with optimized gene and sample leaf orders using Pearson correlation and average linkage clustering. The trees were subsequently cut into clusters using a distance threshold (0.5–1) empirically adjusted to highlight the most relevant features of the trees.

### Comparison of *hrp* cluster sequences

The complete sequences of the *hrp* cluster (*hrpR*–*hrpK*) were retrieved for Psa strains ICMP9617, ICMP19073 and ICMP18884 in the Pseudmonas Genome Database (112) and multiple sequences were aligned using Geneious v11.0.3 (http://www.geneious.com).

### Biofilm assay

Biofilm formation was quantified in Psa1 (J35), Psa2 (KN.2) and Psa3 (CRAFRU8.43) as previously described (113) with slight modifications. Bacterial cells were grown overnight in liquid KB medium before washing and resuspending the cells in KB medium or HIM as described above. The optical density was adjusted to OD_600_ = 0.2 (KB medium) or OD_600_ = 1 (HIM) and 100-μl aliquots of each culture were incubated at 28°C in a transparent 96-multiwell plate for 24 h, shaking at 180 rpm. Cell-free media were used as negative controls. At the end of the incubation period, the microtiter plates were washed with tap water and 150 μl of 0.1% crystal violet was added to each well and incubated at room temperature for 30 min. The plates were rinsed 3–4 times by submerging in water and then blotted carefully on a stack of paper towels to remove excess cells and dye before drying for 30 min. To solubilize the crystal violet, we added 125 μl of 30% acetic acid to each well and incubated at room temperature for 30 min. Absorbance was measured in a plate reader at 550 nm using 30% acetic acid in water as the blank.

### Reverse transcriptase-polymerase chain reaction (RT-PCR)

Total RNA was treated with TURBO DNase (Thermo Fisher Scientific) and first-strand cDNA was synthesized with SuperScript III Reverse Transcriptase (Invitrogen) according to the manufacturer’s instructions. The cDNA was amplified by PCR using GoTaq G2 DNA Polymerase (Promega) with the following parameters: initial denaturation at 95°C for 2 min followed by 30 cycles of denaturation at 95°C for 30 s, annealing at primer-specific temperature (**S2 Table**) for 30 s, and extension at 72°C for primer-specific duration (**S2 Table**), followed by a final fill-in reaction at 72°C for 7 min. Genomic DNA was used as a positive control. The PCR products were separated by 2% agarose gel electrophoresis in 1× TAE buffer. The primer sets are listed in **S2 Table**.

## Acknowledgments

We acknowledge Dr. Marco Scortichini for providing the Psa strains, Prof. Giuseppe Firrao for providing genomic information about strain CRAFRU8.43, Prof. Nicola Vitulo for providing bioinformatics support, and Regione Veneto for funding.

## Supporting information

**S1 Fig. Title**. Legend

**S1 Table.** Title. Legend

## Figure captions

**S1 Appendix. Nucleotide sequences in FASTA format retrieved from the different annotation resources (GenBank, RefSeq, Università di Udine) for the different Psa strains.**

**S2 Appendix. Protein sequences in FASTA format retrieved from the different annotation resources (GenBank, RefSeq, Università di Udine) for the different Psa strains.**

**S3 Appendix. Unique coding sequences (N=20561) used for eArray probe design.**

**S4 Appendix. Complete microarray annotation of 20**,**561 unique coding sequences and their corresponding probes.**

**S5 Appendix. Output of Blast2GO analysis including mapping and annotations of Gene Ontology terms for each probe/unique coding sequence.**

**S6 Appendix. The 17**,**047 unique protein sequences retrieved from unique CDS.**

**S7 Appendix. Complete annotation including protein IDs, microarray probes and superclusters allowing data conversion.**

**S1 Table. Bacterial strains used in this study and genomic annotation resources used for microarray chip design.**

**S2 Table. Primers used in this study.**

**S1 Fig. Flow chart showing the multi-strain custom microarray design procedure.**

**S2 Fig. Principal component analysis based on (A) bacterial strains and (B) growth conditions.** (A) Strains are indicated with different colors (blue = Pto DC3000, purple = V-13, black = CRAFRU 8.43, red = J35, green = KN.2). (B) Growth conditions are indicated in red for rich medium (KB) and green for *hrp*-inducing medium (HIM).

**S3 Fig. Number of non-redundant differentially expressed genes identified in the different Psa strains comparing minimal and rich media at 4 and 8 h post-inoculation.**

**S4 Fig. Common and unique differentially expressed genes (A) upregulated or (B) downregulated in *hrp*-inducing medium compared to KB rich medium in CRAFRU8.43 and V-13 at 4 and 8 h post-inoculation.**

**S5 Fig. Sample clustering based on Gene Ontology terms overrepresented among differentially expressed genes upregulated or downregulated in the different Psa strains cultured in *hrp*-inducing medium for 4 and 8 h.**

**S6 Fig. Differential expression of *hrp*/*hrc* genes in Pto grown in *hrp*-inducing medium compared to rich medium at 4 and 8 h post-inoculation.** Bar length indicates the log_2_ fold change in relative expression of each gene of the *hrp* cluster. Color blocks indicate the different operons of the *hrp* cluster. Arrows on the left indicate operon orientation.

**S7 Fig. Activity of the *hrpA1* promoter in different Psa biovars grown in *hrp*-inducing medium.** Promoter activity was monitored using strains transformed with a reporter system as previously described (114).

**S8 Fig. Expression levels of the main TTSS regulators in different Psa biovars grown in *hrp*-inducing medium compared to KB rich medium.** Bars indicate the log_2_ fold change in relative expression of genes encoding TTSS regulators. Yellow indicates upregulation (log_2_FC > 1) and blue indicates downregulation (log_2_FC < –1).

**S9 Fig. Alignment of the 500-bp sequences upstream of *hrpP* in Psa1, Psa2, Psa3 and Pto.** Nucleotide sequences for Psa1 (ICMP9617), Psa2 (ICMP19073), Psa3 (ICMP18884) and Pto DC3000 were retrieved from the Pseudomonas Genome Database and aligned using Clustal Omega. Nucleotides highlighted in red indicate the *hrp* box consensus motif (GGAACC/T-N_15_-CCAC-N_2_-A).

**S10 Fig. Operon prediction in the *hrp* cluster of Psa3 ICMP18884.** The figure shows the output of the prediction generated by Operon Mapper.

**S11 Fig. Expression of *hrpP, hrpO* and *hrcN* genes in Psa3 grown for 4 and 8 h in KB rich medium and *hrp*-inducing medium (HIM).**

**S12 Fig. Detection of polycistronic *hrpO*-*hrpP* transcripts in Pto DC3000.** Total RNA extracted from cells cultured in rich medium (KB) or *hrp*-inducing medium (HIM) for 4 or 8 h was reverse transcribed and amplified by PCR. Genomic DNA (g) was used as a positive control. The anticipated product sizes were 1747 bp for amplicon *hrcN-hrpO* (primers P1-P2), 890 bp for amplicon *hrpO-hrpP* (primers P3-P4) and 657 bp for the pseudogene transcripts (primers P5-P6).

**S13 Fig. Detection of transcripts encoded by the pseudogene IYO_RS14625 in Psa3 ICMP18884.** Total RNA extracted from Psa1, Psa2 or Psa3 cells cultured in *hrp*-inducing medium (HIM) for 8 h was reverse transcribed and amplified by PCR. Genomic DNA (g) was used as a positive control.

**S14 Fig. Activity of the *hrpA1* promoter in different Psa biovars grown in minimal medium at different temperatures.** Cell were grown in *hrp*-inducing medium (HIM) at 18°C, 24°C or 28°C for 4 h. Promoter activity was evaluated by the quantification of green fluorescent protein used as a reporter.

**S15 Fig. Analysis of the *hrpL* promoter in different Psa biovars.** (A) The 300-bp sequence upstream of *hrpL* in Psa1 (ICMP9617), Psa2 (ICMP19073) and Psa3 (ICMP18884) was retrieved from the Pseudomonas Genome Database and aligned using Clustal Omega. (B) Promoter prediction in the 300-bp sequence upstream of *hrpL* gene in Psa1/Psa2 using BPROM (Softberry) software. (C) Promoter prediction as for (B) bug using Psa3. The transcription factor-binding site (GcvA) identified in Psa3 is highlighted in red.

**S16 Fig. Features of the hybrid histidine kinase (HHK) potentially encoded by the pseudogene strongly expressed in Psa3.** (A) Localization of the pseudogene (IYO_RS14625) in the Psa3 (ICMP18884) genome and the corresponding HHK gene (A262_11517) in Psa2 (ICMP19073), and surrounding genes, including the response regulator (RR) upstream and EAL-containing genes downstream. (B) Alignment of a portion of the HHK-encoding gene (A262_11517) from Psa2 (ICMP19073) and the corresponding pseudogene (IYO_RS15ì4625) from Psa3 (ICMP18884) showing the deletion of five nucleotides in the pseudogene (highlighted in red in the Psa2 sequence). (C) Protein sequence predicted from the translation of Psa3 pseudogene (IYO_RS14625). The figure shows the output of GeneMarkS software (115). (D) Putative protein structure from the translated sequences predicted in (C). Protein domains were analyzed with SMART (116,117).

